# Cysteine Rich Intestinal Protein 2 is a copper-responsive regulator of skeletal muscle differentiation

**DOI:** 10.1101/2024.05.03.592485

**Authors:** Odette Verdejo-Torres, David C. Klein, Lorena Novoa-Aponte, Jaime Carrazco-Carrillo, Denzel Bonilla-Pinto, Antonio Rivera, Fa’alataitaua Fitisemanu, Martha L. Jiménez-González, Lyra Flinn, Aidan T. Pezacki, Antonio Lanzirotti, Luis Antonio Ortiz-Frade, Christopher J. Chang, Juan G. Navea, Crysten Blaby-Haas, Sarah J. Hainer, Teresita Padilla-Benavides

## Abstract

Copper (Cu) is an essential trace element required for respiration, neurotransmitter synthesis, oxidative stress response, and transcriptional regulation. Imbalance in Cu homeostasis can lead to several pathological conditions, affecting neuronal, cognitive, and muscular development. Mechanistically, Cu and Cu-binding proteins (Cu-BPs) have an important but underappreciated role in transcription regulation in mammalian cells. In this context, our lab investigates the contributions of novel Cu-BPs in skeletal muscle differentiation using murine primary myoblasts. Through an unbiased synchrotron X-ray fluorescence-mass spectrometry (XRF/MS) metalloproteomic approach, we identified the murine cysteine rich intestinal protein 2 (mCrip2) in a sample that showed enriched Cu signal, which was isolated from differentiating primary myoblasts derived from mouse satellite cells. Immunolocalization analyses showed that mCrip2 is abundant in both nuclear and cytosolic fractions. Thus, we hypothesized that mCrip2 might have differential roles depending on its cellular localization in the skeletal muscle lineage. mCrip2 is a LIM-family protein with 4 conserved Zn^2+^-binding sites. Homology and phylogenetic analyses showed that mammalian Crip2 possesses histidine residues near two of the Zn^2+^-binding sites (CX2C-HX2C) which are potentially implicated in Cu^+^-binding and competition with Zn^2+^. Biochemical characterization of recombinant human hsCRIP2 revealed a high Cu^+^-binding affinity for two and four Cu^+^ ions and limited redox potential. Functional characterization using CRISPR/Cas9-mediated deletion of *mCrip2* in primary myoblasts did not impact proliferation, but impaired myogenesis by decreasing the expression of differentiation markers, possibly attributed to Cu accumulation. Transcriptome analyses of proliferating and differentiating *mCrip2* KO myoblasts showed alterations in mRNA processing, protein translation, ribosome synthesis, and chromatin organization. CUT&RUN analyses showed that mCrip2 associates with a select set of gene promoters, including *MyoD1* and *metallothioneins*, acting as a novel Cu-responsive or Cu-regulating protein. Our work demonstrates novel regulatory functions of mCrip2 that mediate skeletal muscle differentiation, presenting new features of the Cu-network in myoblasts.

## INTRODUCTION

Copper (Cu) is a trace element crucial for diverse biological processes mediated by Cu-dependent enzymes (1, 2). Examples include cytochrome *c* oxidase (COX) for aerobic respiration, lysyl oxidases (LOXs) for collagen and elastin maturation, Cu-dependent tyrosinase in melanin formation, and autophagy (3–8). Cu also plays essential roles in transcription regulation, catecholamine synthesis, modification of neuro-hypophyseal peptide hormones, methane and ammonia oxidation, and redox homeostasis through superoxide dismutases (SOD1 and SOD3) and ceruloplasmin (CP) (9–16). Recent evidence points to novel functions of Cu in tissue and organ development, including muscle, intestine, and the brain (9, 14, 15, 17–21). Maintaining balanced Cu levels is crucial to prevent the adverse effects of either deficiency and overload, which can lead to various disease states (22, 23). Excessive Cu can cause oxidative stress, initiated by the production of reactive oxygen species (ROS) through the Haber-Weiss and Fenton-like reactions (24). Oxidative stress affects the integrity of multiple molecules, including [Fe–S] cluster proteins and DNA (25). To counter these deleterious effects, the cells employ a finely tuned network of Cu^+^-dependent transcription regulators, soluble chaperones, cuproenzymes, and membrane transporters to uphold intracellular Cu homeostasis and compartmentalization (1, 22, 23, 26–28).

Our understanding of the involvement of Cu-binding proteins (Cu-BP) in transcription regulation in mammalian cells remains limited. Experimental evidence describes the involvement of some metal-binding transcription factors (TFs) and Cu-chaperones, such as the metal regulatory transcription factor 1 (MTF1), specificity protein 1 (SP1), and potentially the antioxidant 1 copper chaperone (ATOX1) in the regulation of gene expression in a Cu-dependent manner (29–33). For instance, MTF1, a highly conserved Zn-binding TF, recognizes and binds to metal-responsive elements (MREs) in genes, maintaining metal and redox homeostasis. MTF1 also regulates the expression of metallothionein genes (*Mt*), and various metal transporters to address metal imbalances (30, 34–39). Beyond its role in metal and redox maintenance, MTF1 is important for development, as demonstrated using murine *Mtf1*-knockout models, flies, and primary myoblasts (9, 40, 41). Studies in *Drosophila* suggested a connection between MTF1 and Parkinson’s disease (PD), where MTF1 overexpression in parkin-mutant flies partially restores the wild type (WT) phenotype, hinting at a potential role in muscle development (42). In this context, our group demonstrated that MTF1 undergoes nuclear translocation to promote the activation of specific myogenic genes once skeletal muscle differentiation has begun. MyoD1, the master regulator of the skeletal muscle lineage, interacts with MTF1 and co-binds at myogenic promoters (9). In line with the significant need for Cu in skeletal muscle differentiation (17), supplementing Cu to differentiating myoblasts enhances Cu^+^-binding to the N-terminus of MTF1, which is essential for the binding of MTF1 and MyoD1 to myogenic genes in differentiating primary myoblasts (9).

Transcription activation operates in a lineage-specific manner. Cell differentiation is orchestrated by master regulators that initiate the expression of specific markers. Skeletal muscle differentiation serves as an exemplary model for unraveling fundamental principles of tissue-specific gene expression and differentiation (43). This system is particularly valuable for investigating the role of Cu, since there is an extensive and well-documented knowledge base on TFs, chromatin remodeling enzymes, and other co-activators and co-repressors. The refined regulation of myoblast determination and differentiation is a tightly regulated process. Initiated by MyoD1, this process is followed by the activation of the proliferation marker Pax7. Subsequently, the activation of Myogenin and other myogenic factors that induce differentiation highlight the refined regulated nature of skeletal muscle tissue (43–69). During myogenesis, metabolic and morphological changes associated with Cu biology manifest, with energy production and redox homeostasis being the most characterized aspects (41, 70, 71). This intrinsic demand for Cu by the skeletal muscle further emphasizes the suitability of this model for studying Cu biology (17).

We proposed that the regulatory roles of Cu during myogenesis extend beyond mitochondrial biogenesis, energy production and maintenance of metal homeostasis. We proposed that Cu regulates a novel category of Cu-BPs that may contribute to the fine-tuned regulation of skeletal muscle differentiation. To address this hypothesis, we employed an unbiased synchrotron-based metalloproteomic approach that identified the murine cysteine rich intestinal protein 2 (mCrip2) as a novel Cu-BP expressed in primary myoblasts derived from mouse satellite cells. Crip2 is a member of the evolutionarily conserved CRIP family, which has been implicated in autophagy (5). Crip2 is expressed and regulated by Wnt3 in migratory cardiac neural crest cells in zebra fish (72). In cutaneous squamous cell carcinoma (cSCC), the interaction between HOXA9 and CRIP2 at gene promoters impedes HIF-1α binding, thus reducing glycolysis (73). Conversely, decreased expression of HOXA9 increases glycolysis, while inhibition of HOXA9 by onco-miR-365 acts as a tumor suppressor by obstructing HIF-1α and its glycolytic regulators. Additionally, evidence suggests that hsCRIP2 interacts with the cuproprotein ATOX1, serving as a negative regulator of autophagy in response to Cu in H1299 cancer cells (5).

Considering these data, we hypothesized that mCrip2 is a Cu-responsive protein, which may regulate skeletal muscle differentiation. Biochemical characterization of purified recombinant hsCRIP2 showed binding of two and four equivalents of Cu per monomer with high affinity. CRISPR/Cas9-mediated deletion of *Crip2* in murine primary myoblasts derived from satellite cells revealed that it is required for differentiation. RNA-seq analyses showed that deletion of *mCrip2* alters the expression of genes involved in mRNA processing, protein translation, ribosome synthesis, and chromatin organization. CUT&RUN and confocal microscopy analyses showed that mCrip2 is a cytosolic and nuclear DNA-binding protein that associates with a limited but specific set of gene promoters in response to Cu-supplementation. This research reveals novel regulatory functions of mCrip2, as a potential Cu-responsive protein that facilitates the transcriptional activation of essential genes for skeletal muscle differentiation and metal regulation. These findings shed light on the intricate regulation of the Cu network within myoblasts, highlighting essential features crucial for proper cellular function and differentiation.

## MATERIALS AND METHODS

### Sequence analyses of the CRIP family

CRIP-like sequences were identified through reciprocal best Blastp searches against the proteomes of the organisms listed in **Supp. Figure 1** using the CRIPs from *Homo sapiens* (HsCRIP1 and HsCRIP2) as the query. The identified hits were then used as queries against the human proteome using the UniProt database (74) or NCBI databases. Only those proteins with HsCRIP1, HsCRIP2, or HsCRIP3 as the top-scoring Blastp hit were kept and used for bioinformatic analyses. Domain identification was performed using InterProScan version 5.65-97.0 (European Bioinformatics Institute, EMBL-EBI, Cambridgeshire, United Kingdom (75)). For phylogenetic reconstruction of individual LIM domains, sequences corresponding to PF00412 plus 20 amino acids upstream and downstream of the domain were retrieved. For all phylogenetic reconstruction, sequences were aligned with MAFFT on XSEDE using CIPRES (76), edited in Jalview (77), and analyzed with IQ-Tree (78). Maximum likelihood trees were constructed using the WAG+I+G4 substitution model and ultrafast bootstrap (1000 replicates). Consensus trees were visualized, rooted at the midpoint, and annotated with iTOL (79). Multiple sequence alignments were visualized with ESPrit 3.0 (80). Data files are available in **Supp. Table 1**. The model structure of murine Crip2 (Af-Q9DCT8-F1-model_v4) was downloaded from the AlphaFold database (EMBL-EBI; (81, 82)). Molecular graphics and annotations were performed with UCSF ChimeraX, developed by the Resource for Biocomputing, Visualization, and Informatics at the University of California, San Francisco, with support from NIH R01-GM129325 and the Office of Cyber Infrastructure and Computational Biology, National Institute of Allergy and Infectious Diseases (83–85).

### Cell culture

Murine primary myoblasts were purchased from iXCells Biotechnologies (10MU-033) and cultured on plates treated with 0.01% Matrigel (354234; Corning, Inc.). Proliferating myoblasts were maintained in growth medium containing 1:1 *v/v* DMEM:F-12 (11039-021; Life Technologies), 20% fetal bovine serum (16140071; FBS, Life Technologies), 25 ng/ml of basic fibroblast growth factor (FGF, 4114-TC; R&D Systems, Inc.), 5% chicken embryo extract (C3999; United States Biological Corporation), and 1% antibiotics (penicillin G/streptomycin, 151-40-122; Gibco) in a humidified atmosphere containing 5% CO_2_ at 37°C. Differentiation was induced by shifting the cells into differentiation media consisting of DMEM (11965-092; Life Technologies), 2% horse serum (16050122; Life Technologies), 1% antibiotics, supplemented or not with a mixture of Insulin/Transferrin/Selenium (51300-044; Gibco) and 30 µM CuSO_4_ or 30 µM tetraethylenepentamine (TEPA) as described (9, 17, 41). Specific culture conditions established for proliferating and differentiating myoblasts are indicated in the figure legends. Briefly, proliferating myoblasts were maintained for 48 h in growth medium supplemented or not with 100 µM CuSO_4_ or 100 µM of the chelator TEPA; differentiating myoblasts were supplemented with 30 µM CuSO_4_ or 30 µM of TEPA, as previously described (9, 17, 41).

HEK293T and BOSC23 cells purchased from ATCC (Manassas, VA) were used to produce lentiviral and retroviral particles, respectively. HEK293T cells were maintained in growth media containing DMEM supplemented with 10% FBS and 1% antibiotics in a humidified incubator at 37°C with 5% CO_2_. GPT media was used to select BOSC23 cells containing the packaging proteins: DMEM, high glucose, no glutamine (11960077, Gibco), 10% FCS, 25X xanthine sodium salt (6.25 mg/ml, X2001, Sigma), 50X HAT supplement (21060017, Gibco) to a final working concentration of 100 μM hypoxanthine, 0.4 μM aminopterin, 16 μM thymidine (Sigma Aldrich H0262), 1% Mycophenolic Acid (MPA; 475913, Sigma), and 1% L-Glutamine (from a 0.2 M stock; A2916801, Gibco). Retroviral particles were collected from BOSC23 maintained in DMEM media supplemented with 10% FBS, 1% antibiotic solution, 1% L-glutamine (from a 0.2 M stock).

### Plasmids construction

To produce recombinant hsCRIP2 protein, human mRNA was isolated from HEK293 cells and used as a template to amplify and clone the gene into the pPRIBA1 vector (IBA), which adds a C-terminal streptavidin (Strep) tag sequence for purification. The primers used are listed in **Supp. Table 2**.

To knockout (KO) the murine *Crip2* (mm*Crip2*) gene in primary myoblasts, we constructed a CRISPR/Cas9 plasmid using three custom-designed sgRNAs to recognize the intron/exon junction 1, 2, and 6 of the *Crip2* murine gene (Reference: ENSMUSG00000006356; **Supp. Table 2**). Each gRNA consisted of 20 nucleotides complementary to the sequence that precedes a 5’-NGG protospacer-adjacent motif (PAM) located in the targeted intron. Specificity was validated by a search through the entire genome to avoid off-target effects. The CRISPR/Cas9 lentiviral constructs were generated using the lentiCRISRv2 oligo cloning protocol (86). Briefly, sense and antisense oligos obtained from Integrated DNA Technology (IDT) were annealed and phosphorylated to form double stranded oligos. These were then cloned into the BsmBI-BsmBI sites downstream from the human U6 promoter of the lentiCRISPRv2 plasmid kind gift from Dr. F. Zhang ((86, 87); Addgene plasmid #52961; Cambridge, MA, USA). The empty vector (EV) encoding for Cas9 but without sgRNA was used as null KO control.

To generate the retroviral constructs to recover expression of WT *hsCRIP2*, the coding sequence was PCR amplified from the pPRIBA vector used for expression of hsCRIP2 in bacteria. PCR products were subsequently cloned into the pBABE retroviral vector containing a blasticidin resistance cassette (88) using the primers listed in **Supp. Table 2**. All constructs were confirmed by sequencing.

### Virus production, and transduction of primary myoblasts

To generate lentiviral particles, 5×10^6^ HEK293T cells were plated in 10 cm dishes. The next day, cells were transfected with 15 µg of the murine *Crip2* sgRNA-containing CRISPR/Cas9 vector mixed with the packaging plasmids pLP1 (15 µg), pLP2 (6 µg) and pSVGV (3 µg). Retroviruses were produced by transfecting 15 µg of the pBABE construct encoding the human *CRIP2* gene in BOSC23 cells (93). Transfections were performed using Lipofectamine 2000 (Invitrogen) following the manufacturer’s instructions. The following day, the media was replaced with 10 ml of fresh DMEM supplemented with 10% FBS. The viral supernatant was harvested after 24 and 48 h and filtered through a 0.22 µm syringe filter (Millipore). To infect primary myoblasts, 5 ml of the filtered supernatant supplemented with 8 μg/ml polybrene (Sigma) were used to transduce 2 × 10^6^ cells, as previously described (9, 41). After overnight incubation, infected cells were selected in growth media containing 2 μg/ml puromycin (Invitrogen). Stable myoblasts were maintained in growth media containing 1 μg/ml puromycin.

### Antibodies

The primary rabbit anti-Crip2, -strep-tag, -Gapdh and -vinculin antibodies were obtained from ABclonal (A9038, AE066, A19056 and A2752, respectively; Woburn, MA, USA). The mouse anti-metallothionein was obtained from GeneTex (GTX12228, Irvine, CA, USA); and anti-Lamin A/C and anti-normal IgG were from Santa Cruz Biotechnology Inc (sc-376248, sc-2025, respectively, Dallas, TX, USA). The rabbit anti-Tubulin was obtained from Cell Signaling Technology (2144, Danvers, MA, USA). Hybridoma supernatants against Pax7 (deposited by A. Kawakami), myosin heavy chain (MF20, deposited by D. A. Fischman) and myogenin (F5D, deposited by W. E. Wright) were obtained as hybridoma supernatants from the Developmental Studies Hybridoma Bank (The University of Iowa, Iowa City, IA, USA). The secondary antibodies used were goat anti-rabbit and -mouse coupled to HRP and the fluorescent goat anti-rabbit Alexa-633 and -mouse Alexa-488 (31460, 31430 and A21070, A11001, respectively; Thermo Fisher Scientific).

### Protein expression and purification

The pPRIBA plasmid encoding *hsCRIP2* was transformed into *Escherichia coli* BL21, (DE3) competent cells. Protein expression was performed following the autoinducing media protocol (89). Purification of CRIP2 protein was performed using Strep-Tactin resin (2-12-01-025; IBA) for soluble cuproproteins, following the manufacturer’s instructions. Purified proteins were stored at -20°C in buffer containing 10% glycerol, 100 mM Tris, pH 8, and 150 mM NaCl. Protein concentrations were determined using the Bradford assay (90). Molar protein concentrations were estimated assuming that the proteins were pure, using MW = 22,500. To eliminate any bound metal, purified proteins were treated with metal chelators as previously described (9, 91–94). Briefly, the proteins were incubated for 45 min at room temperature with 0.5 mM EDTA and 0.5 mM tetrathiomolybdate. Chelators were removed by buffer exchange using 10 kDa cut-off Centricons (UFC801096; Millipore Sigma). The final purity of all protein preparations was ∼95%, as verified by Coomassie Brilliant Blue staining of SDS-PAGE and western blot.

### Metal loading to recombinant purified proteins

Cu^+^ and Zn^2+^-loading to apo-hsCRIP2 was performed as previously described (91, 93) with minor modifications. Cu^+^- and Zn^2+^-loaded hsCRIP2 was obtained by incubating the protein (10 µM) with a 10-fold molar excess of CuSO_4_ or ZnSO_4_ in 25 mM Tris-HCl (pH 8.0), 50 mM NaCl, 10 mM ascorbic acid for 15 min at room temperature with gentle agitation. The unbound metals were removed by centrifuging in a 10-kDa cut-off Centricon (Millipore Sigma) and 5 volumes of deionized water for washing. A fraction of the metal-loaded samples was used to measure protein content using Bradford reaction (90). The remaining metal loaded samples were mineralized with excess of fuming HNO_3_ (trace metal grade) for 1 h at 80 °C and 2M H_2_O_2_ for 60 min at room temperature. Cu and Zn content was measured by furnace atomic absorption spectroscopy (AAS, Perkin Elmer).

### Cu^+^-binding affinity determination for purified hsCRIP2

The dissociation constant (KD) of hsCRIP2 for Cu^+^ was determined through competition assays with the chromogenic ligand bathocuproine disulfonate BCS ([Cu^+^(BCS)_2_]^3−^ β_2_’ formation constant 10^20.8^ M^-2^, ε_483 nm_ 13000 M^-1^ cm^-1^) (95) as previously shown (96). Immediately before the assay, CRIP2 was fully reduced by incubation with 5 mM TCEP on ice for 3 h, and the reducing agent was removed using a 3 kDa Centricon (Millipore Sigma). Cu^+^ solutions were generated in situ from CuSO_4_ in the presence of ascorbate. Briefly, 10 µM Cu^+^, 25 µM BCS in buffer 25 mM HEPES pH 8, 150 mM NaCl, 10 mM ascorbic acid were titrated with 1 - 30 µM purified CRIP2, incubated 5 min at room temperature, and the 300-800 nm absorption spectra were recorded. CRIP2-Cu^+^ K_D_ was calculated by curve-fitting the experimental data to the equilibrium equations [1] and [2] (97), where M is the metal, L is the competing ligand and P are the Cu-sites in the protein. Reported errors are asymptotic standard errors provided by the fitting software (KaleidaGraph; Synergy).

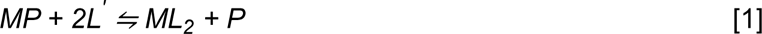

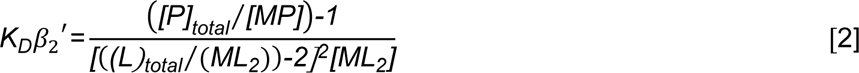

### Cyclic voltammetry

Recombinant purified human CRIP2 protein was metallated by mixing with Cu(NO_3_)_2_ in a 1:1 molar ratio with mild agitation for 24 h at room temperature. *Electrode cleaning:* A polycrystalline gold electrode (ɸ=3 mm) was cleaned by mechanical polishing with diamond powder and 0.3 μm alumina; then it was electrochemically treated in a 0.5 M solution of H_2_SO_4_ cycling from -0.25 to -1.125 V vs Hg/HgSO_4_ using a scan rate of 5 mVs-1 until the characteristic gold profile was observed. *Formation of the self-assembled monolayer of L-Cysteine (L-Cys)*: The treated electrode was incubated with a 5 mM solution of L-Cys in 0.1 M phosphate buffer pH 7.2. Then, the electrode was washed with deionized water to remove the excess thiol. *Electrochemical characterization:* Measurements were performed at room temperature (25 °C) in a 2 mL cell using a potentiostat-galvanostat Biologic SP-300 (Seyssinet-Pariset, France) with 0.1 M phosphate buffer pH 7.2 as supporting electrolyte in the absence and in the presence of protein (50 µM). A typical three-electrode arrangement was used. Modified and not modified gold electrode were used as working electrodes, a platinum electrode as a counter electrode and an Ag/AgCl electrode as reference. All solutions were bubbled with nitrogen prior each experiment. During measurements, a nitrogen atmosphere was maintained in the electrochemical cell. The cyclic voltammograms were obtained with a potential window of -500 to 500 mV at scan a rate range of 10 to 2500 mVs-1.

### Metalloproteomic analyses

Analysis of cuproproteins in differentiating murine myoblasts was performed by 2-dimension grazing-exit X-ray fluorescence (2D-GE/XRF) coupled to liquid chromatography-mass spectrometry (LC-MS/MS) as previously described (41, 98, 99). Briefly, 100 μg of whole cell protein extract from primary myoblasts stimulated to differentiate in the presence or absence of insulin or Cu were extracted 24 h post-stimulation. Samples were resolved on 10% native polyacrylamide gels to preserve the metal co-factors bound to the proteins. Gels were blotted onto PVDF membranes and analyzed using synchrotron micro-XRF at the GSECARS beamline 13-ID-E, at the Advanced Photon Source (APS), Argonne National Lab (Illinois, United States) as previously described (41). Briefly, the incident X-ray beam energy was tuned to 10.2 keV for these analyses and focused to a relatively broad spot size of ∼150 × 200 mm using rhodium-coated silicon mirrors in a Kirkpatrick-Baez geometry (100). Energy dispersive X-ray fluorescence spectra were collected using a four-element silicon drift detector (Vortex ME4, SII NanoTechnology). Spectra were collected in mapping mode (20 mm × 70 mm maps) by raster scanning the beam through the incident beam in a continuous scan mode, resulting in maps with a 250 mm pixel size and an accumulation time of 80 ms per pixel. Maps of total measured Cu Kα fluorescence intensity were generated and normalized to incident flux, using the GSECARS Larch software (101). The PVDF membranes were marked along the blot periphery, and the micro-XRF mapping images were overlaid onto the membranes. Subsequently, bands aligning with the Cu signals were excised for tryptic digestion and subsequent identification via LC-MS/MS at the Proteomics and Mass Spectrometry Facility of the University of Massachusetts Chan Medical School. The data obtained was analyzed using Scaffold_3.5.1 software from Proteome Software Inc.

### Primary myoblast immunofluorescence

Primary myoblasts were cultured on CellView Advanced TC culture dishes (Grenier Bio-One, Monroe, NC, USA). Myoblasts grown under proliferation conditions were harvested after 48 h of seeding, and differentiating cells were fixed 24 h after induction of differentiation. The cells were fixed in 10% formalin, permeabilized with PBT buffer containing 0.5% Triton X-100 in PBS, and then blocked in 5% horse serum in PBT for 1 h. Next, the samples were incubated with a rabbit anti-Crip2 antibody (1:100) in blocking solution overnight at 4°C, washed 3 times with PBT for 10 min at room temperature and incubated with the corresponding secondary fluorescent antibodies (1:100; Thermo Fisher Scientific) in blocking solution for 2 h at room temperature. Nuclei were stained for 30 min with DAPI. Samples were mounted with VectaShield solution (Vector Laboratories, Newark, CA) and imaged with a Leica SP8 using the 63X water immersion lens. Images were analyzed with the Leica Application Suite X (Leica Microsystem Inc).

### Immunohistochemistry and fusion index determination

Proliferating and differentiating primary myoblasts were grown in 48-well plates and fixed with 10% formalin in PBS overnight at 4°C and permeabilized with 0.2% Triton X-100 in PBS for 20 min at room temperature. Immunohistochemistry (IHC) against Pax7, myogenin, and myosin heavy chain was performed using hybridoma supernatants for these antigens and the Universal ABC Kit. IHCs were developed using the Vectastain Elite ABC HRP Kit (both from Vector Laboratories) following the manufacturer’s instructions. Images were acquired with an Echo Rebel microscope using the 20X objective (Echo, San Diego, CA).

Three independent biological replicates were used to calculate the fusion index, which was determined as the ratio of the number of nuclei in myocytes with 2 or more nuclei *vs.* the total number of nuclei. Images were analyzed using ImageJ software v.1.8 (National Institutes of Health; (102)).

### Western blot analysis

Proliferating and differentiating primary myoblasts were solubilized in RIPA buffer [10 mM piperazine-N,N-bis(2-ethanesulfonic acid) pH 7.4, 150 mM NaCl, 2 mM ethylenediamine-tetraacetic acid (EDTA), 1% Triton X-100, 0.5% sodium deoxycholate, and 10% glycerol] supplemented with Complete protease inhibitor cocktail (Thermo Fisher Scientific). The lysates were sonicated for 6 min, in cycles of 30 s on-off at mild intensity using a Bioruptor UCD-200 (Diagenode, Denville, NJ). Protein concentrations were quantified using the Bradford method (90) and 20 µg of protein were separated on 10% SDS-PAGE and transferred to PVDF membranes. Primary antibodies were used at 1:1000 dilutions and incubated overnight at 4°C with agitation. The following day, the membranes were incubated with species-specific secondary antibodies for 2 h and treated with horseradish peroxidase (HRP) substrate for enhanced chemiluminescence (ECL; Tanon, AbClonal Technologies). Densitometric analyses were performed with ImageJ software v.1.8 (National Institute of Health; (102)).

### Cell proliferation assays

Three independent biological replicates of control and *Crip2* KO primary myoblasts were initially seeded at a density of 1×10^4^ cells/cm^2^ in proliferation media supplemented or not with 100 µM CuSO_4_. Sampling and cell counting were performed at 24, 48, and 72 and 96 h post-plating. The cells were subjected to trypsinization, followed by triple washing with PBS, and subsequently counted using a Spectrum Cellometer from Nexcelcom Biosciences (Lawrence, MA).

### CUT&RUN

CUT&RUN was performed as described (103–107), using recombinant Protein A-MNase (pA-MNase;(108)). Briefly, nuclear extraction was performed on 100,000 cells with a hypotonic buffer (20 mM HEPES-KOH, pH 7.9, 10 mM KCl, 0.5mM spermidine, 0.1% Triton X-100, 20% glycerol, fresh protease inhibitors) and bound to lectin-coated concanavalin A magnetic beads (40 µL beads per 100,000 nuclei; Polysciences, Warrington, PA). Bead-bound nuclei were chelated with blocking buffer (20 mM HEPES, pH 7.5, 150 mM NaCl, 0.5mM spermidine, 0.1% BSA, 2mM EDTA, fresh protease inhibitors) and washed (wash buffer: 20 mM HEPES, pH 7.5, 150 mM NaCl, 0.5mM spermidine, 0.1% BSA, fresh protease inhibitors). Nuclei were incubated in wash buffer containing primary antibody (anti-CRIP2, Abclonal A9038, lot 0050480101; rabbit polyclonal IgG, Abcam ab37415, lot #GR3208186-1) for 1 h, followed by 30 min incubation in wash buffer containing pA-MNase. All antibody and pA-MNase incubations were carried out at room temperature with rotation. Samples were equilibrated to 0°C in an ice-water bath and 3 mM CaCl_2_ was added to activate pA-MNase. After 22 min, the digestion was inactivated with 20 mM EDTA and 4 mM EGTA, and 1.5 pg MNase-digested *Sacharomyces cerevisiae* mononucleosomes were added as a spike-in control. Genomic fragments were released after an RNase A treatment and subsequent centrifugation. Isolated fragments were used as input for a library build consisting of end repair and adenylation, ligation of NEBNext stem-loop adapters, and purification with AMPure XP beads (Agencourt). Barcoded fragments were then amplified by 15 cycles of high-fidelity PCR and purified using AMPure XP beads. Libraries were pooled and sequenced on an Illumina NextSeq2000 to a depth of ∼10 million uniquely mapped reads.

### CUT&RUN data analysis

Paired-end fastq files were trimmed to 25 bp and mapped to the mm10 genome with bowtie2 (options -q -N 1 -X 1000; (109)). Duplicate reads were identified and removed with Picard (110) and reads were filtered for mapping quality (MAPQ ≥ 10) with SAMtools (111). Size classes corresponding to factor-bound footprints (1-120 bp) were generated using SAMtools and a custom awk script (111). Reads were converted to bigWig files using deepTools with RPKM normalization (options -bs 1 --normalizeUsing RPKM; (112)). CUT&RUN peaks were called using the CUT&RUN-specific peak calling program SEACR (version 1.3, options: non, 0.0001; (113)). Heatmaps were plotted using deepTools computeMatrix (options -a 2000 -b 2000 -bs 20 -- missingDataAsZero) and plotHeatmap (112). Gene Ontology (GO) analysis was performed on peaks present in both CUT&RUN replicates using Metascape, with the y-axis representing rank of enrichment (114, 115).

### Chromatin immunoprecipitation (ChIP)

Three independent biological replicates of differentiating primary myoblasts were crosslinked with 1% formaldehyde for 10 min at room temperature with continuous mild shaking. Fixation was deactivated using glycine, and cells were further incubated for 5 min at room temperature. Subsequently, samples were washed three times with 10 ml of ice-cold PBS supplemented with Complete Protease Inhibitor (Roche, Basel, Switzerland), and the cells were resuspended in 1 ml of the same buffer. After centrifugation for 5 minutes at 5,000 × g at 4°C, the PBS was discarded, and primary myoblasts were lysed using the SimpleChIP Plus Sonication Chromatin IP Kit (56383; Cell Signaling Technology) following the manufacturer’s instructions. Cells were sonicated using a Bioruptor UCD-200 (Diagenode) for six cycles of 30 s on and 30 s off at mild intensity. The samples were then incubated with the anti-Crip2 antibody or IgG as a negative control. Immunoprecipitated material was collected using magnetic beads and washed with 1x ChIP buffer supplemented with NaCl according to the manufacturer’s recommendations. Subsequently, samples were eluted in 1x elution buffer, incubated overnight at 65°C, and subjected to reverse crosslinking with NaCl. The resulting DNA was purified using the ChIP DNA clean concentrator as per the manufacturer’s instructions (Zymo Research, Irvine, CA, United States) and stored at -80°C until further analysis by (qRT-PCR). The primer sequences in the promoter regions of *MyoD1, Mt1*, and *Mt2* are listed in **Supp. Table 2**.

### RNA-Seq

RNA-seq was performed as previously described (107). Control myoblasts transduced with the CRISPR/Cas9 EV and KO cells transduced with *Crip2* sgRNA-3 were cultured as detailed above. RNA was extracted with TRIzol per manufacturer’s instructions and purified by chloroform extraction and isopropanol precipitation. The purified RNA was aliquoted for sequencing and RT-qPCR analyses, flash-frozen in liquid nitrogen and stored at -80 °C until use. Ribosomal RNA was depleted from 2 µg of input RNA via antisense tiling oligonucleotides and digestion with thermostable RNase H (MCLabs) (116). rRNA-depleted RNA samples were treated with Turbo DNase (ThermoFisher) and purified by silica column (Zymo RNA Clean & Concentrator kit). Strand-specific RNA-seq libraries were built from rRNA-depleted RNA, using the NEBNext Ultra II Directional Library kit. RNA was fragmented at 94 °C for 15 min and subsequently used as input for cDNA synthesis and strand-specific library building, per manufacturer’s protocol. Libraries were amplified for 8 cycles of high-fidelity PCR, then pooled and sequenced to a depth of approximately 20 million uniquely mapped reads on an Illumina NextSeq2000.

### RNA-seq data analysis

Paired-end fastq files were aligned to the mm10 mouse genome using STAR (options -- outSAMtype BAM SortedByCoordinate --outFilterMismatchNoverReadLmax 0.02 -- outFilterMultimapNmax 1). To visualize RNA-seq data, bigwigs were generated using deepTools with TPM read normalization (options -bs 5 –smoothLength 20 --normalizeUsing BPM; (112)). Feature counts were generated using subread featureCounts (options -s 2 -p -B)(117). Count files were read into R and downstream analysis was conducted using DESeq2 with the apeglm log2 fold change shrinkage correction applied (118, 119). Differentially expressed genes were plotted using EnhancedVolcano (120). Gene ontology (GO) analysis was performed on significantly up- and downregulated genes separately using Metascape (115). Gene lists used for GO analyses contained only significantly altered genes with a DESeq2 baseMean value of ≥ 1 to remove lowly expressed genes. Significance was defined as DESeq2 adjusted p-value < 0.05. Y-axes indicate pathway enrichment ranking.

### Quantitative reverse transcription PCR

Aliquots of samples obtained for RNA-seq of proliferating and differentiating primary myoblasts were used as templates for cDNA synthesis (1 µg) and quantitative reverse transcription (qRT-PCR). Briefly, the RNA was treated with HiFiScript gDNA Removal RT MasterMix Kit (CW2020M, Cwbio, Jiangsu, China), following the manufacturer’s protocol. Quantitative PCR was performed with FastSYBR Mixture (CW0955L, Cwbio) on the AriaMx Real-Time PCR System (Agilent) using the primers listed in **Supp. Table 2**. The delta threshold cycle value (2^ΔΔCT^; (121)) was calculated for each gene and represented the difference between the CT value of the gene of interest and that of the control gene, *Eef1a1α*.

### Integration of CUT&RUN and RNA-seq datasets

CUT&RUN and RNA-seq datasets were integrated based on gene lists that were identified as significantly up- or downregulated per DESeq2 results (padj. < 0.05) and CUT&RUN peak files as generated and described above. A bed file containing differentially expressed gene (DEG) promoters was generated using the UCSC Table Browser (122), with promoters defined as regions spanning 1 kb upstream of UCSC RefGene transcription start sites. Direct overlaps between CUT&RUN peaks and DEG promoters were assigned using the HOMER mergePeaks function (options -d given; (114)). Overlapping regions were annotated using the HOMER annotatePeaks.pl function and the resulting file was manually separated by specific overlap groups (114). Bed files for each combination of CUT&RUN peaks and DEG promoter regions were generated and used as input for subsequent Gene Ontology, Genome Ontology, and motif enrichment analyses via the HOMER annotatePeaks.pl and findMotifs.pl functions (114). A background gene list of all genes expressed in these experiments (DESeq baseMean value ≥ 1) was used to control for potential overrepresented motif sequences.

### Metal Content Analysis

Three independent biological replicates of control and *Crip2* KO primary myoblasts were cultured under proliferating and differentiating conditions as described above. Cells were fractionated using the Rapid, Efficient, and Practical (REAP) nuclear and cytoplasmic separation method (123). Briefly, myoblasts were washed with ice-cold PBS, scraped, and transferred to a 1.5 ml microcentrifuge tube. After centrifugation for 10 s at 13,000 x *g*, the supernatant was discarded. The cell pellet was resuspended in 500 µl of ice-cold PBS containing 0.1% NP-40 (Millipore Sigma), and 100 µl of the cell suspension were collected as the whole cell fraction. The remaining 400 µl were utilized to obtain nuclear and cytosolic fractions by disrupting the cells through pipetting using a 1 ml pipette tip. After another 10 s of centrifugation, the supernatant was collected as the cytosolic fraction. The nuclear pellet was washed twice in 1 ml of ice-cold PBS containing 0.1% NP40, followed by an additional 10 s of centrifugation. The supernatant was removed, and the pellet was resuspended in 100 µl of PBS. Nuclear integrity was confirmed using light microscopy. Subsequently, all samples underwent sonication using a Bioruptor at medium intensity for 5 min with 30 s on-off cycles. Protein quantification was performed using the Bradford method (90). The purity of the fractions was assessed by western blot analysis.

Comparative analysis of metal concentrations was carried out by triplicate measurements of Cu in each sample, using AAS equipped with a graphite furnace (GF-AAS; PerkinElmer, PinAAcle 900T) with Cu hollow cathode lamp as radiation source as previously described (9, 41, 124–127). This technique enables ultra-trace elemental analysis (<1 ppm) in low volume samples, where dilution can be restricted due to the low initial concentration of analyte in the sample. A known mass of nuclei and cytosolic fractions were acid digested in concentrated HNO_3_, using a single-stage digestion method. To prevent contamination during ultra-trace analysis, all reagents for sample treatment and calibration standard preparation were of analytical grade and diluted in purified water with a resistivity of 18 MΩ. All analytical glassware and materials were thoroughly cleaned with a 3% HCl solution for 24 h, and an ultra-clean graphite furnace was employed for all analyses. Cu standard solutions (1000 mg L^-1^; Sigma-Aldrich) were diluted as necessary to obtain working standards and determine the limits of detection and dynamic range of the method. The limit of detection (LOD) for Cu, calculated as three times the standard deviation (3σ), was determined to be 0.01 ppb, with a limit of linearity (LOL) at 160 ppb. Cu content on each sample was normalized to the initial cell mass.

### Visualization of labile copper in primary myoblasts using fluorescent sensors

To visualize labile Cu-pools in proliferating and differentiating myoblasts we used three different membrane-permeable fluorescent dyes CS1, CD649 and CD649.2 following standard procedures reported with minor modifications (128–130). Briefly, independent biological replicates of control and *Crip2* KO myoblasts were seeded in CellView Plates and cultured in the presence or absence of CuSO_4_ and insulin as indicated in the figure legends. Prior to incubation with the Cu probes, the cells were washed with PBS supplemented with 1X GlutaMax (Thermo Fisher Scientific) and 10 mM CaCl_2_. The cells were imaged immediately after incubation with the probes using a Leica SP8 confocal microscope. The images were captured and analyzed with the Leica Application Suite X (Leica Microsystem Inc, Deerfield, IL).

The CS1 probe visualized Cu^+^ under 543 nm excitation and 566 nm emission; the CD649 probe, which detects Cu^+^ and Cu^2+^, and the CD649.2 probe for Cu^2+^ were imaged at 633 nm excitation, and fluorescence emission was collected in the range of 650-750 nm (128–130). Cells were washed with PBS, then incubated with both probes (5 µM) in the dark for 10 min. After washing, cells were imaged immediately using a Leica SP8 microscope and analyzed with Leica Application Suite X.

### Statistical analyses

In all cases, the data represents the mean of three independent biological replicates ± SE. Statistical analyses were performed using Graph Pad Prism 7.0b (Dotmatrics, Boston, MA). Multiple data point comparisons and statistical significance were determined using one-way analysis of variance (ANOVA) followed by Bonferroni multiple comparison tests. Experiments where p< 0.05 were considered statistically significant.

### Data availability

Genomic data sets have been deposited in the Gene Expression Omnibus (GEO) accession no. GSE252162.

## RESULTS

### Unbiased identification of Cu-binding to mCrip2 in primary myoblasts via XRF/MS analysis

We have previously reported that Cu is essential for proliferation and differentiation of primary myoblasts, as depleting the culture medium of the ion inhibits these processes (9, 17, 41). We established a primary myoblast culture model for inducing differentiation by withdrawing growth factors and introducing non-toxic concentrations of CuSO_4_ (30 µM), resulting in phenotypes comparable to those seen in control cells cultured with insulin (131, 132). To better understand the role of Cu in differentiation and identify proteins that connect Cu availability to myoblast development, we identified Cu-binding proteins from different stages of myoblast development using an unbiased X-ray fluorescence-mass spectrometry (XRF/MS) approach (**Fig. 1A**). Whole cell extracts of differentiating myoblasts treated with or without insulin or Cu were separated by non-denaturing PAGE and transferred into PVDF membranes to preserve Cu-protein interactions. The samples were then analyzed by XRF-MS, revealing a Cu signal in the middle of the membrane for the myoblast sample differentiated with Cu (**Fig. 1A**, indicated by the red box). Mass spectrometry analyses of this Cu-enriched fraction showed mCrip2 as the most relevant hit for Cu^+^-binding due to its Cys and His signature pattern, and subsequently identified as a cuproprotein that interacts with ATOX1 and modulates autophagic response in the lung cancer cell line H1299 (5).

**Figure 1.**
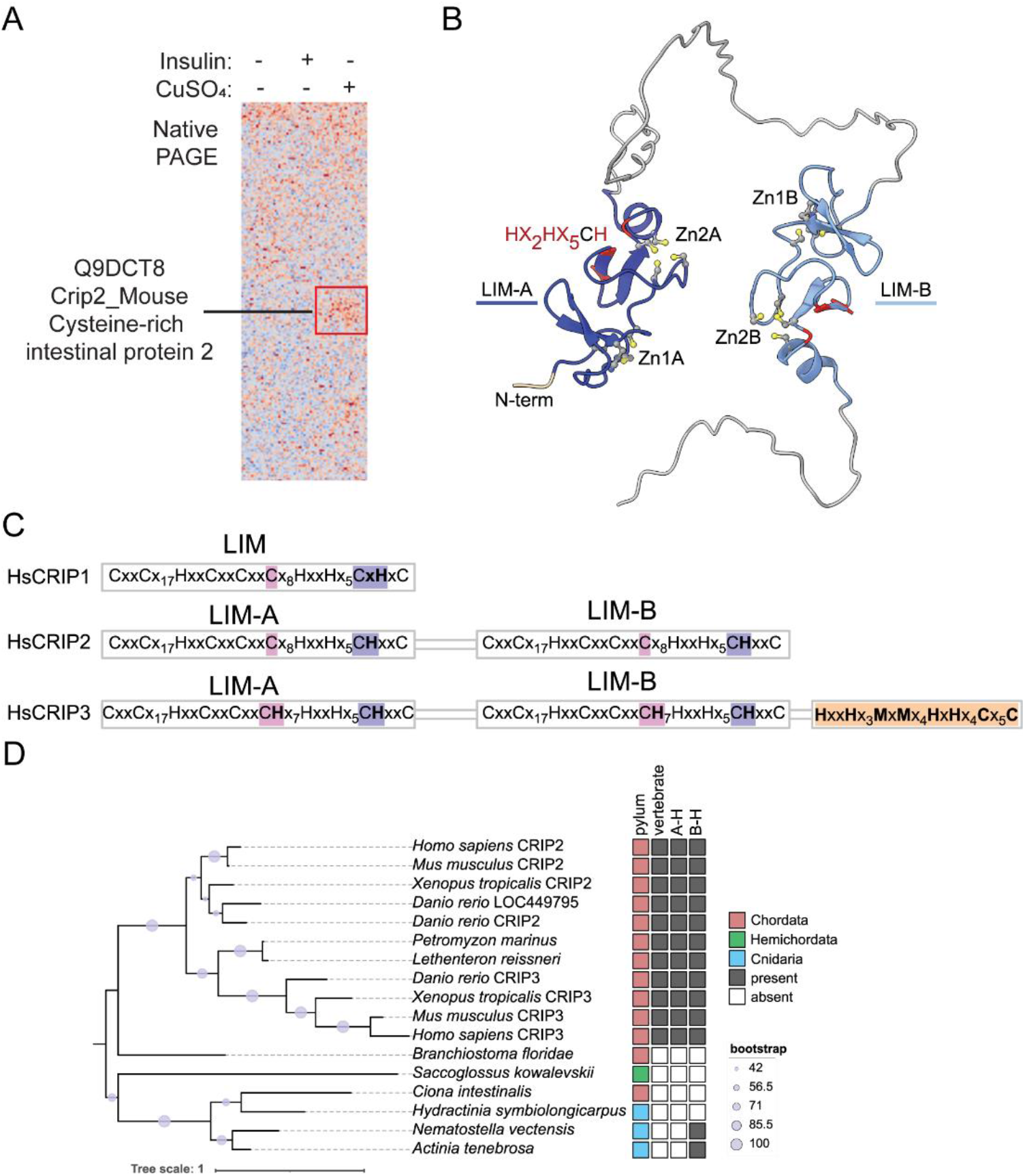
Phylogenetic analyses of the CRIP family members and appearance of the Cu^+^-binding sites. **A.** Synchrotron based X-Ray fluorescence analyses coupled to protein sequencing by mass spectrometry of native PAGE gels from whole cell extracts of differentiating (24 h) primary myoblasts showed the presence of mCrip2 in a section that showed enriched signal of copper (indicated by red box). **B.** AlphaFold-predicted structure of mCrip2. The N-terminal LIM domain is colored dark blue and the C-terminal LIM domain is colored light blue. The conserved Zn-binding residues are shown as balls and sticks, while the conserved histidine residues located near the Zn2 sites are colored red. **C.** Metal-binding motifs found in human CRIPs. Individual LIM domains are shown as boxes and labeled. Individual Zn-finger motifs are shaded with grey and labeled. The CRIP-specific HxxX motif is highlighted in yellow. In addition to having duplicated LIM domains, CRIP2 and CRIP3 proteins contain conserved histidine residue adjacent to Cys53 and Cys172 (purple shading). Human CRIP3 contains a histidine residue next to Cys35 in the Zn2 sites that are not found in CRIP1 or CRIP2 (pink shading). **D.** Maximum likelihood tree of CRIPs with 2 two LIM domains. “A-H” refers to conservation of human CRIP2 His36 and “B-H” refers to conservation of human CRIP2 His173.

### Phylogenetic analysis of CRIP2

The CRIP family, named after the Cysteine-rich intestinal protein (CRIP;(133)), is comprised of three paralogs encoded with the mouse and human genomes: CRIP1, CRIP2, and CRIP3. hsCRIP1 contains a single LIM domain (named for Lin11, Isl-1, and Mec-3; defined by InterPro: PF00412), whereas hsCRIP2 and hsCRIP3 contain two LIM domains separated by an amino acid stretch that is predicted to be largely unstructured (**Fig. 1B**). Here the N-terminal domain is denoted as “LIM-A” and the C-terminal domain as “LIM-B”. Like other characterized LIM domains, each is anticipated to adopt a double Zn-finger fold with the CxxC···HxxC motif, referred as Zn1, and the CxxC···CxxC motif referred as Zn2 (**Fig. 1C**). Based on structural characterization of rat Crip1 (134) and the LIM-A domain of hsCRIP2 (PDB: 2CU8), and supported by conservation of Zn^2+^-binding residues found in other LIM family proteins, hsCRIP2 and hsCRIP3 are expected to be able to bind four Zn^2+^ ions (**Fig. 1B**). Diverging from closely related LIM-containing proteins, human and mouse CRIPs contain conserved histidine motifs (e.g., Hx2Hx5CH in hsCRIP2, where the C is the third Cys residue of the Zn2 site) near the Zn2 sites (**Fig. 1B,C**), while hsCRIP3 also contains a C-terminal region rich in putative metal-binding motifs.

Phylogenetic reconstruction and motif analysis of the two-LIM domain CRIPs suggests that Crip2/Crip3 are specific to vertebrates (**Fig. 1D**). Although homologous proteins found in non-vertebrates and Cnidaria contain the Crip-specific HxxH motif, they are missing one or both of the histidine residues adjacent to the third cys residue of the Zn2 site (**Fig. 1D, Supp. Fig. 1**). Phylogenetic reconstruction of individual CRIP LIM domains from various Holozoa lineages suggests that the ancestral vertebrate Crip was likely similar to CRIP1 with a duplication event subsequently leading to the emergence of the two-domain Crip2/Crip3 (**Supp. Fig.1**). This analysis supports the existence of the two-domain CRIP in the common ancestor of vertebrates based on presence in lampreys that are from one of the earliest diverging lineages of vertebrates compared to mammals. The analysis also suggests that the two-domain Crips could have been present earlier in Deuterostomia evolution with the presence of two-domain Crips in a lamprey and acorn worm. However, in the case of these invertebrates, it is not clear whether the domain expansion occurred independently from the domain expansion leading to Crip2/Crip3 in vertebrates. For example, the *Branchiostoma floridae* genome is predicted to encode a two-LIM domain Crip, while the *B. lanceolatum* genome is predicted to encode two single-LIM domain proteins (**Supp. Fig. 2**). Since the *B. lanceolatum* proteins are encoded by side-by-side genes, it is possible that either the *B. floridae* or *B. lanceolatum* gene models are inaccurate, and the LIM domains should be separate proteins in *B. floridae* or that the separate proteins in *B. lanceolatum* should be a single fusion protein (**Supp. Fig. 2**). Two-LIM domain CRIPs were not identified in Bilateria outside of Deuterostomia. There is a conserved protein containing a N-terminal LIM domain and a C-terminal Ras-associated domain orthologous to mammalian Rassf, which is found throughout the animal lineage but absent in Deuterostomia (**Supp. Fig. 2**). Bootstrap support for the phylogenetic relationships involving the two-domain cnidarian CRIPs is too weak to speculate on the timing of their appearance (**Supp. Fig. 2**). However, the occurrence of closely related one-domain and two-domain CRIPs suggests the possibility of independent duplication events resulting either in two-domain CRIPs or independent events of domain loss.

### Biochemical characterization of hsCRIP2 as a high-affinity Cu^+^-binding, redox-neutral protein

Based on the presence of conserved His residues unique to vertebrate CRIPs and their proximity to two of the Zn-binding sites, we postulated that Cu^+^ could bind at or near the Zn sites and induce a conformational change. To address this hypothesis, we cloned, expressed, and purified the Strep-tagged WT hsCRIP2. Representative Coomassie brilliant blue staining and western blots against CRIP2 and Strep-tag show the purity of the protein at a molecular weight of approximately 22 KDa. A band detected around 40 KDa may indicate the presence of dimers in the preparation (**Fig. 2A**). We determined the Cu^+^-binding properties of hsCRIP2 by incubating the purified, chelated protein with 10-fold molar excess of CuSO_4_ in the presence of ascorbate as a reducing agent. After removing the unbound metal, the protein was mineralized, and stoichiometry of binding was determined by atomic absorbance spectroscopy (AAS). hsCRIP2 can bind 3.71 ± 0.37 Cu^+^ atoms per protein, 4.15 ± 0.47 atoms of Zn, and approximately two Cu and two Zn ions when simultaneously incubated with both metals (**Table 1**). No Cu^2+^ was found to be bound to hsCRIP2 *in vitro*.

**Figure 2.**
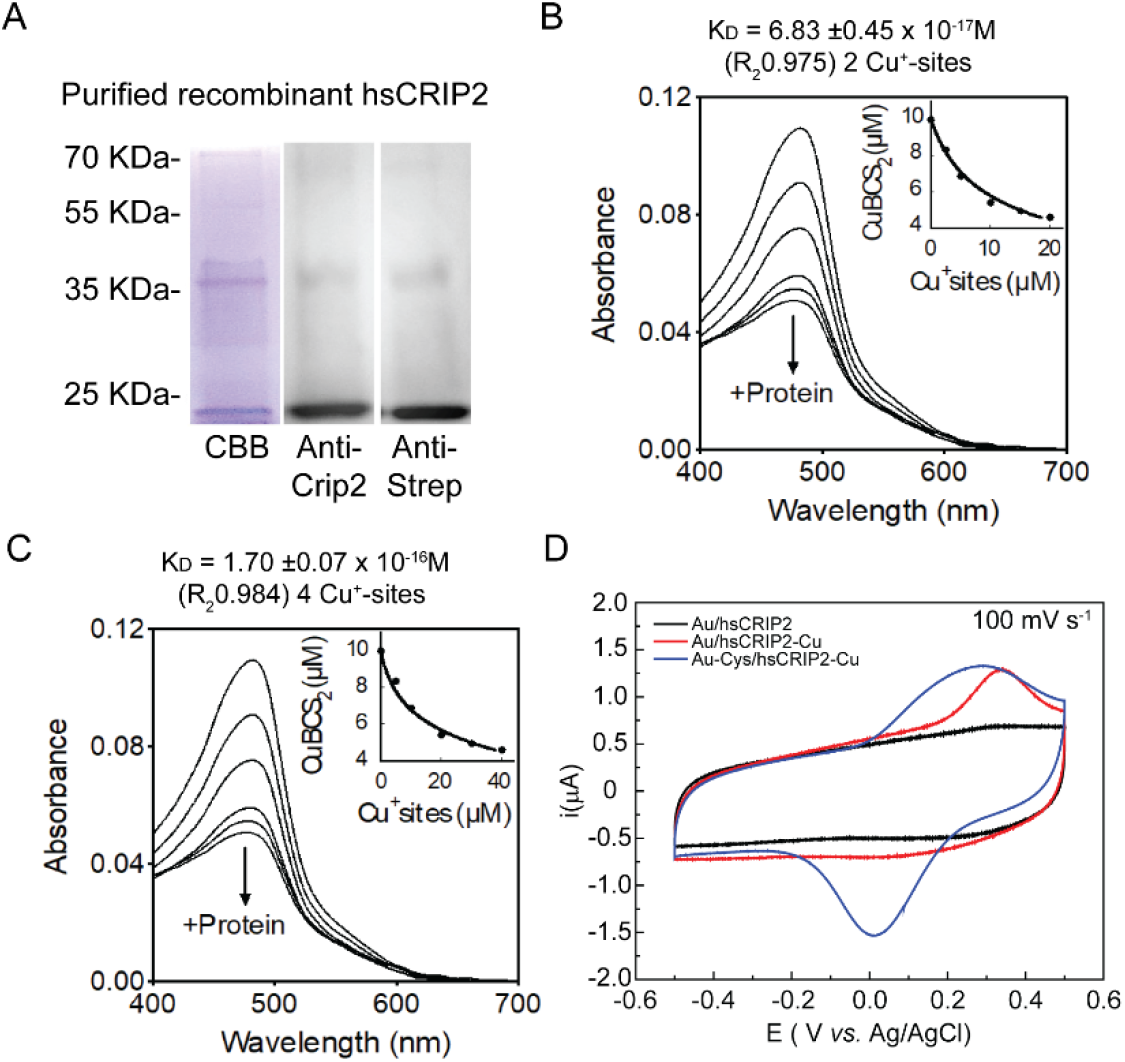
Biochemical characterization of hsCRIP2 as a Cu^+^-binding protein. **A.** Recombinant human hsCRIP2 protein purification; representative Coomassie Brilliant Blue (CBB) gel and western blots of hsCRIP2 protein detected using an anti-Crip2 and anti-Strep-tag antibodies. Determination of the Cu^+^ dissociation constant (*K_D_*) of hsCRIP2 using the BCS competition assay; spectrophotometric titration of 25 µM BCS and 10 μM Cu^+^ with increasing concentrations of hsCRIP2. The arrow indicates the decrease in absorbance at 483 nm upon each protein addition. hsCRIP2-Cu^+^ dissociation constants were calculated based on the stoichiometry determinations, assuming two Cu^+^ sites **(B**) or four Cu^+^ sites **(C**) per hsCRIP2 molecule. **D.** Cyclic Voltammograms obtained for a) Au/CRIP2, b) Au/CRIP2-Cu and c) Au-Cys/CRIP2-Cu, measured in a potential window of -500 to 500 mV using a scan rate of 100 mV/s with 0.1 M phosphate buffer pH 7.2 as supporting electrolyte.

**Table 1.** Cu+ and Zn^2+^-binding stoichiometry of purified hsCRIP2.

| **Metal** | **hsCRIP2** |
| --- | --- |
| Cu<sup>+</sup> | 3.72 +/- 0.37 |
| Zn<sup>2+</sup> | 4.15 +/- 0.47 |
| Cu<sup>+</sup><br>/Zn<sup>2+</sup> | 2.07 +/- 0.52<br>2.22 +/- 0.29 |

To explore the strength of the Cu^+^-binding capacity of purified hsCRIP2, its Cu^+^-binding affinity was estimated via metal competition experiments using the high-affinity ligand bathocuproine disulfonate (BCS; **Fig. 2B,C**). Excess ligand was added to the reactions to ensure effective competition. hsCRIP2 avidly competed with BCS for Cu^+^ as evidenced by the measurable decrease in the absorbance of the [Cu^+^(BCS)_2_^3-^] complex at 483 nm. The spectral determinations after titrating increasing amounts of hsCRIP2 in the competition experiments fit well to the equilibrium equations (1) and (2), allowing us to determine the corresponding K_D_ values (**Fig. 2B,C**). The metal-to-hsCRIP2 stoichiometry determinations demonstrated that the protein binds 4 Cu^+^ ions in the absence of Zn, however the Cu^+^ occupancy of hsCRIP2 drops to 2 Cu^+^ ions when Zn^2+^ is present (i.e., a 2:1 Zn:hsCRIP2 ratio was observed), which is in agreement with the stoichiometry determinations. The impossibility of discriminating between metal-binding sites during the metal/ligand competition experiments with the current setup, led us to consider two possible mathematical determinations of the K_D_ values: hsCRIP2 binds Cu^+^ either at two or at four thermodynamically indistinguishable metal-binding sites. A dissociation constant K_D_ = 6.83 ± 0.45 × 10^-17^ M was estimated when considering two Cu^+^-binding sites on hsCRIP2. This binding affinity drops to K_D_ = 1.70 ± 0.07 × 10^-16^ M when considering four Cu^+^-binding sites on hsCRIP2. Although the experimental data fits to the equilibrium equation [2] is better when considering 4 Cu^+^-binding sites on hsCRIP2 (**Fig. 2C**), this difference is not significant enough to discriminate between the two proposed scenarios. The estimated K_D_ values for Cu^+^-binding to hsCRIP2 are within the range of affinities observed for many other Cu^+^-binding proteins (97, 135–137) and agree with the proposed virtual absence of free Cu^+^ in the cytosol, including recent determinations of hsCRIP2 (5). Finally, equivalent experiments to determine the Cu^+^-binding affinity of hsCRIP2 in the presence of Zn^2+^ were unsuccessful due to protein precipitation observed when hsCRIP2 was added to a solution containing 25 µM BCS, 10 µM Cu^+^ and 10 µM Zn^2+^. This precipitation was observed at all the tested protein concentrations (1-30 µM; **Supp. Fig. 3A**).

To determine whether the presence of Cu^+^-bound to CRIP2 conferred redox capabilities to the protein, we performed cyclic voltammetry assays, as previously described (138). Briefly, to enhance electron transfer, the electrode surface was modified using gold alkanethiols self-assembled monolayers (SAMs; (139–142)), as these favor protein stability and prevent enzyme denaturation over the electrode surface (143, 144). SAMs also possess low reliance on the electrode surface and superior electron transfer rates (139). In addition, alkanethiols, such as L-Cys, exhibit a robust interaction with the gold surface and possess terminal groups with high affinity for the enzymatic structure, facilitating the electron transfer process, and contribute to the stable immobilization of an enzyme on the electrode surface without affecting catalytic activity (139, 141, 142). To investigate parameters related to the electron transfer processes of hsCRIP2 metallated with Cu^+^, the protein was mounted over an electrode modified with SAM and in the presence and absence of L-Cys. The electron transfer rate constant (k_s_), symmetry coefficient (α), and redox potential parameters for hsCRIP2 were determined as previously reported (138, 141). SOD cyclic voltammetry determinations were used as comparison of an efficient redox active enzyme (138). Cyclic voltammograms obtained for a bare Au electrode in the presence of a solutions of hsCRIP2 and hsCRIP2-Cu showed that for Au/hsCRIP2 system, no redox processes are observed (**Fig. 2D**). Conversely, when the protein is metallated (Au/hsCRIP2-Cu) an oxidation peak at E_pa_ 325 mV vs Ag/AgCl related to the electron transfer process Cu^+^ → Cu^2+^ + 1e^-^ is observed, but no reduction peak corresponding to the reaction Cu^2+^ + 1e^-^ → Cu^+^ was detected (**Fig. 2D**). The above can be attributed to conformational changes in the protein structure that buries the metal center into the protein domain, causing an irreversible electrochemical response (138). It has been reported that the formation of thiol films over gold electrodes improves the electron transfer processes of redox proteins (139). Therefore, the electrochemical response for an Au electrode modified with the L-Cys film (Au/Cys-hsCRIP2-Cu, **Fig. 2D**) resulted in an oxidation peak at ∼275 mV *vs* Ag/AgCl and a reduction peak at 15 mV *vs* Ag/AgCl on the reverse scan. These peaks can be attributed to redox couple Cu^+^/Cu^2+^, with a redox potential value (E°) of 151 mV *vs.* Ag/AgCl and a difference between potential peaks (ΔEp) of 295 mV. Thermodynamic and kinetic parameters related to the electron transfer process were obtained from cyclic voltammograms at different scan rates (10 to 2500 mVs-1) for Au-Cys/hsCRIP2-Cu (**Supp. Fig. 3B**). The linear relationship between anodic and cathodic currents with scan rate (*v*) indicated that the protein is attached to the electrode surface (**Supp. Fig. 3C**). The surface excess concentration (Γ°) and symmetry factor (α) for electron transfer were calculated from I_pa_ *vs. v* and I_pc_ *vs. v* plots. Furthermore, the heterogeneous electron rate constant (k_s_) was obtained by the Laviron method (**Supp. Table 3**). For the L-Cys-hsCRIP2-Cu the values for Γ_o_, α and k_s_ were 1.07×10^-10^ mol cm^-2^, 0.53 and 1.02 s^-1^ respectively. For Au/hsCRIP2-Cu, no reduction signals were observed, even when the scan rate was increased **(Supp. Fig. 3D**); thus, it was not possible to calculate the values of Γ°, α and k_s_. A comparison with values obtained for Cu-SOD, with a Cys Au electrode in the same experimental conditions taken from literature, indicated similar values of symmetry factor for heterogeneous electron rate constant (k_s_), related to the fact that L-Cys promotes electron transfer in the same way, approaching and fixing the redox center near to the electrode surface (**Supp. Table 3**). On the other hand, the different values in redox potential values (E°) surface excess concentration (Γ_o_), indicate a different coordination sphere for the Cu center and different surface protein charge for both proteins that modified the protein concentration at the electrode surface (138). Altogether, the data suggest that Cu-bound hsCRIP2 has limited redox activity.

### Cu supplementation increases the cytosolic expression of mCrip2

To explore the functional connections between Cu and mCrip2 in primary myoblasts derived from mouse satellite cells, we cultured the cells under varying concentrations of the metal. Then, we assessed changes in mCrip2 expression levels through western blot and densitometric analyses (**Fig. 3A,B**). Primary myoblasts cultured in differentiation medium, with or without insulin supplementation, exhibited comparable mCrip2 levels. Yet, differentiation in medium supplemented with 30 µM CuSO_4_ significantly elevated mCrip2 levels. Conversely, chelating Cu with TEPA, which impedes differentiation (17), reduced mCrip2 expression (**Fig. 3A,B**). These findings strongly suggest a role for Cu in inducing mCrip2 during myogenesis. Confocal microscopy images showed that mCrip2 is found in both the nucleus and in the cytosol and presents a punctuated pattern (**Fig. 3C,D**). Upon Cu supplementation, the levels of mCrip2 increased, consistent with the findings from western blot analyses, and became predominantly cytosolic (**Fig. 3**). These observations suggest that Cu prompts the cytosolic accumulation of mCrip2.

**Figure 3.**
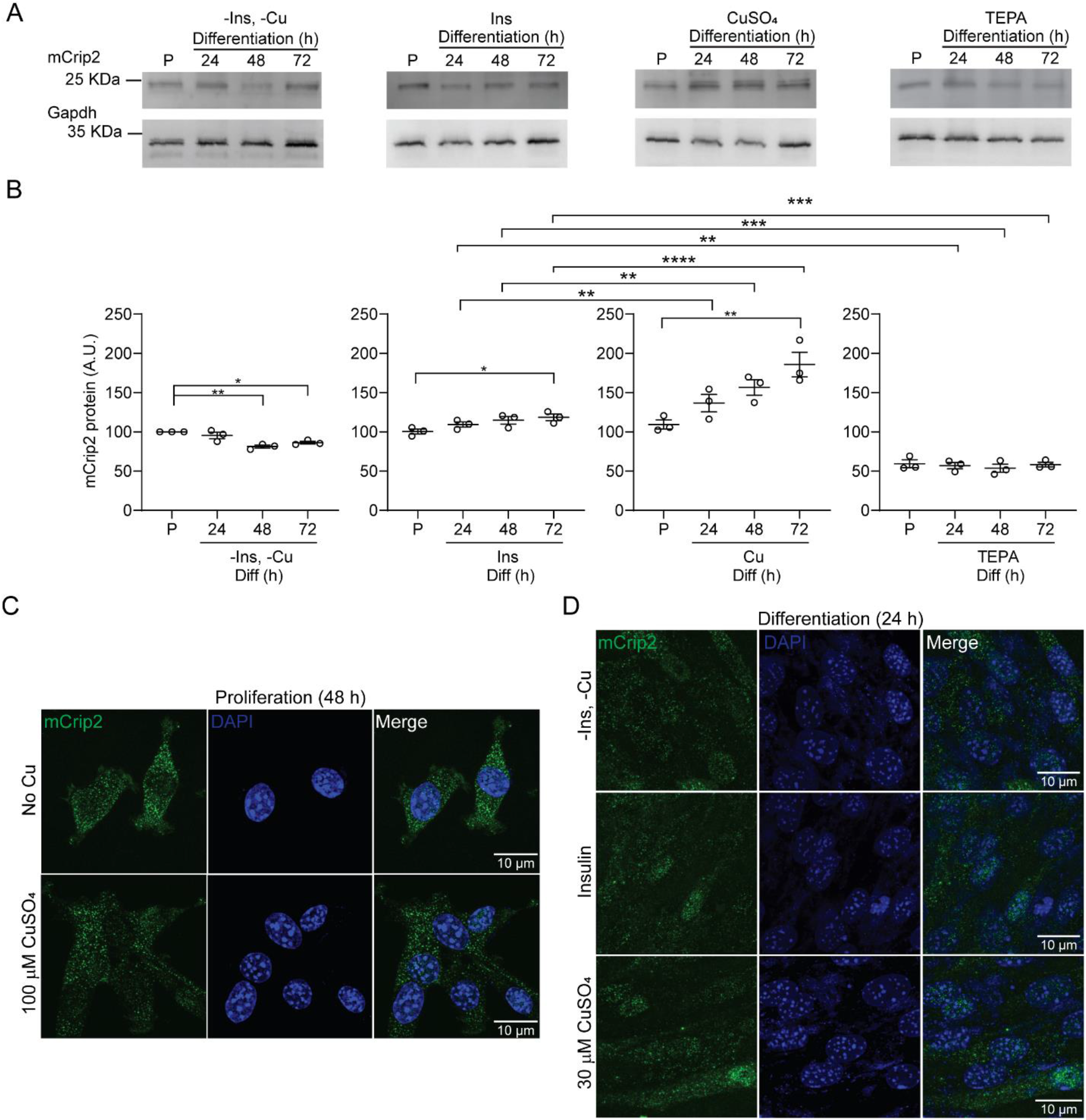
Effect of Cu supplementation in mCrip2 expression and localization in proliferating and differentiating primary myoblasts. **A. B.** Representative western blots of mCrip2 protein levels in proliferating (48 h) and differentiating (24, 48, 72 h) myoblasts in the presence or absence of insulin, CuSO_4_, and the Cu-chelator TEPA. Anti-Gapdh antibody was used as a loading control. **C.** Densitometric quantification of mCrip2 bands in proliferating and differentiating primary myoblasts shown in A. N=5, *P < 0.05; **P < 0.01. Representative confocal microscopy images showing the cytosolic and nuclear localization of mCrip2 (GREEN) in proliferating **(D**) and differentiating **(E**) primary myoblasts. Nuclei were counterstained with DAPI (blue). N=6.

### mCrip2 is required for myoblast differentiation

To elucidate the physiological role of mCrip2 in the skeletal muscle lineage, we employed viral vectors encoding the CRISPR/Cas9 system to generate *mCrip2*-deficient primary myoblasts. Three lentiviral constructs encoding three different sgRNAs targeting intron-exon junctions of *mCrip2* (**Fig. 4A**) were employed to stably knockout (KO) the endogenous protein. Myoblasts transduced with the empty vector (EV)-containing lentivirus served as negative controls. The infected cells were selected with puromycin, and mCrip2 levels were assessed via western blot (**Fig. 4B**) and densitometric analyses (**Fig. 4C**). Proliferating and differentiating myoblasts transduced with either *mCrip2* sgRNAs exhibited a significant decrease in the protein expression compared to WT and EV controls (**Fig. 4B,C; Supp. Fig. 4A**).

**Figure 4.**
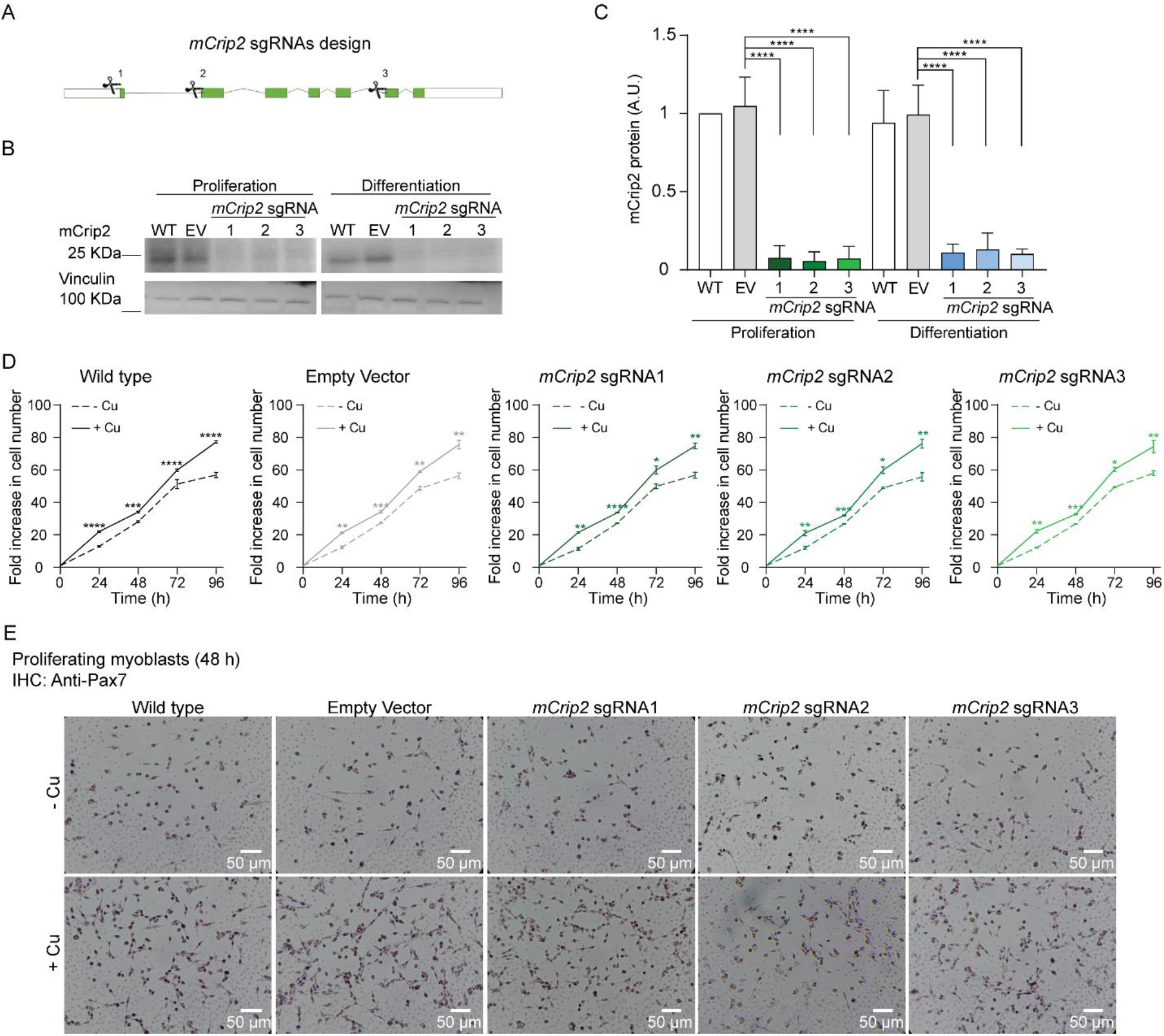
CRISPR/Cas9-mediated knock-out of *mCrip2* in primary myoblasts. **(A)** Location of the three sgRNA designed to delete *mCrip2* by CRISPR/Cas9 system in primary myoblasts. The three constructs were targeted to intron/exon boundaries. **(B**) Representative western blot (top) and **(C**) densitometric quantification (bottom) of mCrip2 protein levels in proliferating (48 h) and differentiating (24 h) primary myoblasts. An anti-vinculin antibody was used as a loading control. N=6; ****P < 0.0001. **(D**) CRISPR/Cas9-mediated deletion of *mCrip2* does not impact proliferation of primary myoblasts. Proliferation curves comparing wild type, empty vector (EV) control, and *mCrip2* depleted (sgRNA1, sgRNA2 and sgRNA3) primary myoblasts. No significant differences in growth were found between the five strains. The Cu-dependent increase in proliferation rate previously reported (Vest et al., 2018) is maintained in *mCrip2* KO myoblasts. Data represent the mean of 3 biologically independent experiments +/-SE. Statistical analyses compared each strain grown in the presence (100 µM) *vs*. absence of CuSO_4_ in the culture media. *P < 0.05; **P < 0.01; ***P < 0.001; ****P < 0.0001. **(E**) Representative light micrographies of proliferating myoblasts immunostained for Pax7 at 48 h cultured with or without 100 µM CuSO_4_; N=3.

First, we assessed the functional effect of *mCrip2* deletion in proliferating myoblasts. Cell counting assays showed that *mCrip2* KO cells exhibited growth kinetics comparable to WT cells, indicating that proliferating primary myoblasts can withstand the depletion of this gene (**Fig. 4D**). In fact, the *Crip2* KO cells presented an enhanced proliferative phenotype that results from Cu supplementation, observed previously in C2C12 cells and primary myoblasts (9, 17). Immunohistochemistry analyses of control and *mCrip2* KO myoblasts also demonstrated that the expression of the proliferation marker Pax7 is not altered by this deletion (**Fig. 4E**). These data indicate that deletion of *mCrip2* had no effects on primary myoblast proliferation.

To assess whether KO of *mCrip2* impairs myogenesis, control and *mCrip2* sgRNA-transduced primary myoblasts were differentiated under varying concentrations of insulin and Cu as previously reported (9, 17, 41). Briefly, the conditions used include a control condition that renders poor differentiation phenotypes by culturing the cells in the absence of insulin (-Ins, -Cu), a positive control of standard differentiation with insulin (+Ins), and a Cu-supplemented (+Cu) culture, which we have shown rescues the absence of insulin from the media leading to differentiation (17). Differentiated myoblasts under these three conditions were fixed at 24 and 48h and stained against myogenic markers to assess the differentiation phenotype **(Fig. 5; Supp. Fig. 4**). Immunohistochemistry analyses using an anti-myogenin or an anti-myosin heavy chain (MHC) hybridoma confirmed that *Crip2* KO cells fail to differentiate. A decreased incidence of myogenin-positive nuclei, formation of myosin-containing myotubes, and decreased fusion index was observed in KO cells compared to WT and EV control myoblasts (**Fig. 5A,B; Supp. Fig. 4,5A**). Microscopy analyses also showed complete detachment of *Crip2* KO cells upon induction of differentiation in the absence of insulin and Cu after 48 h (-Cu, -Ins; **Fig. 5A; Supp. Fig. 4,5A**). Myoblasts differentiated with insulin also detached, and surviving cells showed an approximately 70% decrease in fusion index compared with controls (+Ins, **Fig. 5A,B; Supp. Fig. 4**). Importantly, Cu-supplementation partially prevented cell detachment; however, the cells failed to express the differentiation markers and exhibited a significantly lower fusion index than WT cells (+Cu, **Fig. 5A,B; Supp. Fig. 4,5A**). This data suggests that although Cu partially prevented cell death, the metal was unable to rescue the differentiation defect.

**Figure 5.**
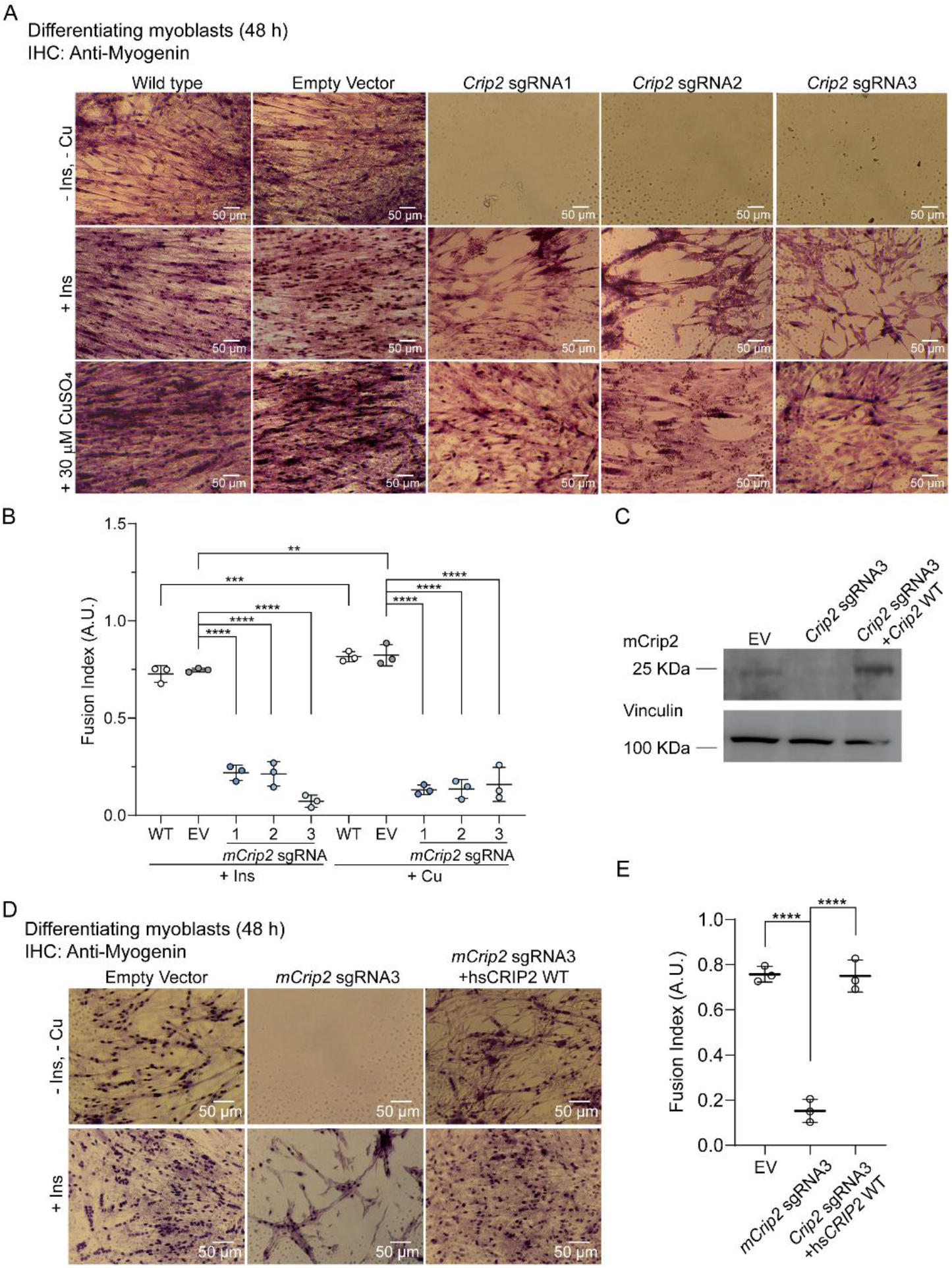
CRISPR/Cas9-mediated KO of *mCrip2* impairs differentiation of cultured primary myoblasts. **(A)** Representative light micrographs of differentiating myoblasts immunostained for myogenin after 48 h of inducing differentiation in the presence or absence of insulin and Cu. **(B**) Calculated fusion index for controls and three different clones of myoblasts transduced with *mCrip2* sgRNAs. Data represent three independent biological experiments +/- SE. N=3; **P < 0.01; ***P < 0.001; ****P < 0.0001. **(C**) The differentiation defect of *mCrip2* KO myoblasts is rescued by reintroducing *hsCRIP2* gene. Representative western blot of primary myoblasts transduced with either empty vector (EV), the *mCrip2* sgRNA3 and the KO *mCrip2* cells transduced with *hsCRIP2* construct. Samples were obtained for 48 h after inducing differentiation. A specific anti-Crip2 antibody depicts the levels of protein and vinculin was used as loading control. **(D**) Representative light micrographs of differentiating myoblasts immunostained for myogenin **(E**) Calculated fusion index for EV control, KO cells and recovered cells with wild type hsCRIP2. Data represents 3 independent biological experiments +/- SE. ****P < 0.0001.

Aiming to rescue the differentiation defect observed for the *mCrip2* KO cells, we reintroduced wild type hs*CRIP2* into the sgRNA3 myoblasts, using a retroviral overexpression system. Western blot analyses showed successful expression of hsCRIP2 in the KO cells (**Fig. 5C**). Then, we tested whether the exogenous protein could rescue the differentiation defect by immunohistochemistry. To this end, EV control cells, the *mCrip2* sgRNA3 clone, and the cells transduced with the retroviral vector were differentiated in the presence or absence of insulin for 48 h, which was the time point where we detected the largest defect (**Fig. 5D**). Microscopy analyses of the differentiation markers myogenin (**Fig. 5D**), MHC (**Supp. Fig. 5B**), and fusion index quantification (**Fig. 5E**) showed that the exogenous hsCRIP2 rescued the differentiation defect. Collectively, these data highlight the essential role of mCrip2 during the differentiation of primary myoblasts derived from mouse satellite cells.

### Primary myoblasts lacking *Crip2* present elevated levels of Cu

Our lab has previously reported that Cu levels increase as a consequence of differentiation, aligning with the inherent necessity of this metal during myogenesis (17). Subcellular fractionation analyses showed an increase of Cu in the nucleus and cytosol during myogenesis (9). However, it is well established that Cu imbalances can lead to deleterious effects (22, 26, 145, 146). Considering the phenotypes observed in the KO *mCrip2* cells, we asked whether normal Cu trafficking was affected by *mCrip2* deletion. Subcellular fractionation of proliferating (**Fig. 6A**) and differentiating myoblasts supplemented with insulin (**Fig. 6B**) revealed that *mCrip2* KO results in elevated levels of Cu in the nuclear and cytosolic fractions for both stages. Complementation experiments using the retroviral overexpression system to reintroduce the WT *hsCRIP2* gene to the KO cells restored the levels of Cu in both proliferating and differentiating myoblasts (**Fig. 6A,B**). Purity of the fractions was confirmed through western blot analysis of lysates from primary myoblasts differentiated with insulin, utilizing lamin A/C to identify the nuclear fraction and β-tubulin for the cytosolic fraction (**Fig. 6C**; (9, 125)).

**Figure 6.**
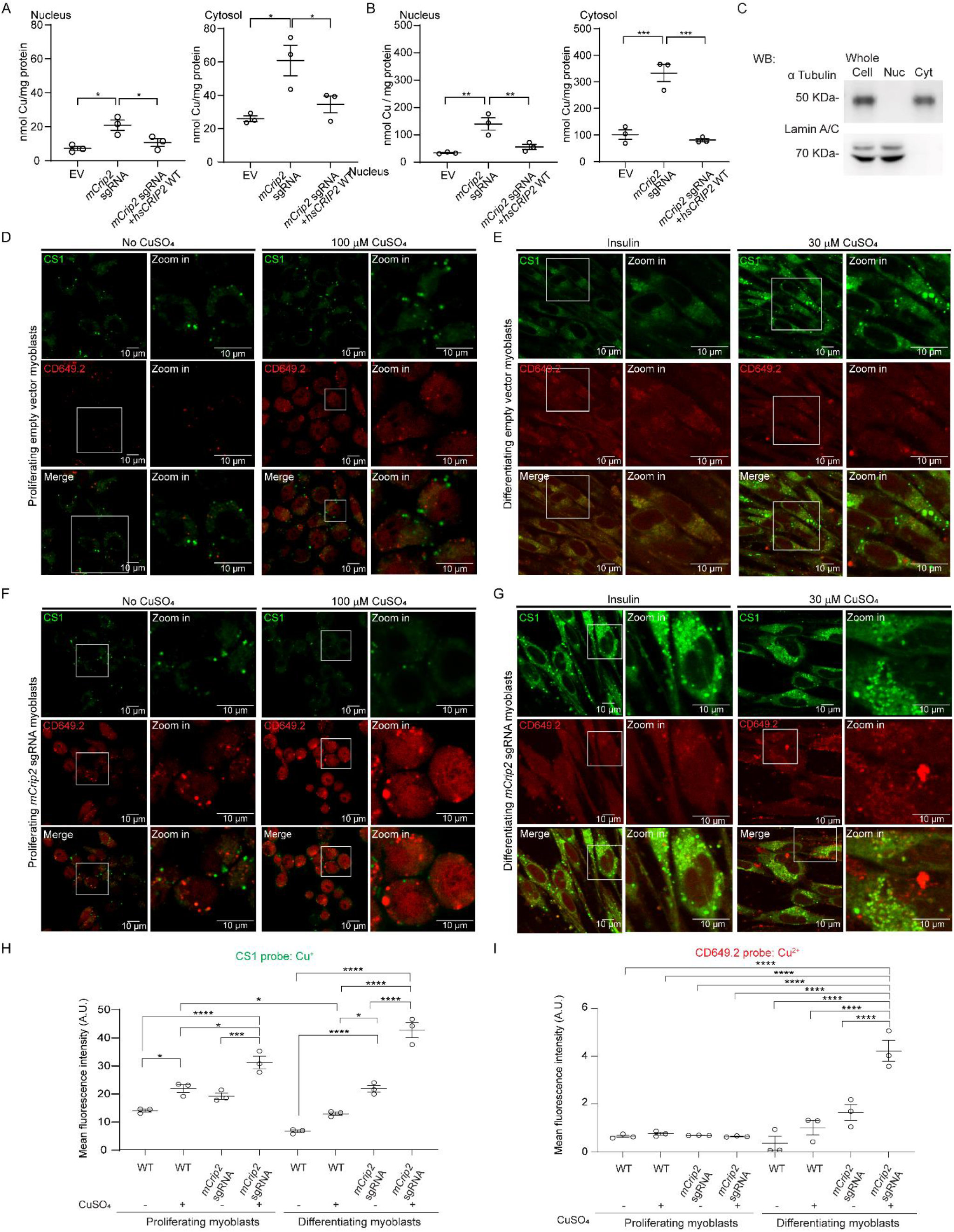
Changes in the cellular distribution of Cu in myoblasts depleted of *mCrip2*. Nuclear and cytosolic Cu content of proliferating **(A**) differentiating **(B**) primary myoblasts determined by AAS. Data represent the distribution of the data obtained from three independent biological experiments; *P < 0.05; **P < 0.01; ***P < 0.001. **(C**) Representative western blot showing the purity of the subcellular fractions. Lamin A/C, and α-Tubulin were used as controls to show the separation of nuclear and cytoplasmic fractions. Myoblasts depleted of *mCrip2* present elevated levels of labile Cu^+^ and Cu^2+^ pools. Live-cell confocal microscopy images of wild type primary myoblasts and KO for *mCrip2* supplemented or not with CuSO_4_. Cu^+^ imaging was performed by incubating the cells with 5 µM CS1 (Cu^+^, green) and CD649.2 (Cu^2+^, red) for 10 min at 25°C. Wild type proliferating **(D**) and differentiating myoblasts **(E**). Primary myoblasts with CRISPR/Cas9-mediated deletion of m*Crip2* under proliferation **(F**) and differentiating conditions **(G**). Quantification of the fluorescence of live-cell imaging for Cu^+^ with the CS1 probe **(H**) and Cu^2+^ with the CD649.2 probe **(I**) in proliferating myoblasts and differentiating myoblasts. N=3, *P < 0.05; ***P < 0.001; ****P < 0.0001.

To determine whether these changes in Cu levels were impacting the labile Cu pools, we performed live cell imaging with three fluorescent probes. These probes can selectively distinguish the metal and its oxidation states, allowing the detection of changes in labile intracellular Cu^+/2+^ pools (128–130). The CS1 probe detects Cu^+^, the CD649 probe senses Cu^+/2+^, and the CD649.2 probe detects Cu^2+^. Myoblasts were cultured in the presence and absence of insulin and CuSO_4_ as indicated in the figure legend and incubated with different combinations of probes (**Fig. 6D-G; Supp. Fig. 6**). Consistent with the AAS data shown here and previously reported (9, 17), we detected increased Cu levels in differentiating cells compared to proliferating myoblasts. Live cell confocal microscopy imaging showed that in both cases monovalent Cu seems to be more abundant and appears to be confined to subcellular compartments or vesicles (**Fig. 6D-G; Supp. Fig. 6**; CS1 probe). When control cells were supplemented with non-toxic concentrations of Cu (100 µM for proliferation and 30 µM for differentiation; (9, 17, 41)) the levels of labile Cu^+^ were slightly elevated versus untreated control myoblasts. Analysis of proliferating and differentiating *Crip2* KO myoblasts showed a significant increase in labile Cu^+^ when cells are cultured with Cu, indicating that the ion that accumulates in the mutant (**Fig. 6D-H; Supp. Fig. 6**; CS1 probe) compared to the EV controls. This data aligns with the biochemical properties of the protein and the observed phenotype in differentiating *mCrip2* KO myoblasts (**Figs. 2, 5**). Given its high Cu K_D_, mCrip2 likely sequesters labile Cu. Since its deletion primarily impacts differentiating myoblasts, detecting higher labile Cu upon knockout is anticipated. We detected minimal changes for labile Cu^2+^ in proliferating myoblasts (**Fig. 6D,F,H**, CD649.2 probe), which slightly increased in proliferating and differentiating cells cultured with Cu (**Fig. 6E,G,I**, CD649.2 probe). These data were validated with the CD649 probe which may detect both Cu^+^ and Cu^2+^ (**Supp. Fig. 6**; CD649 probe). Considering the significant accumulation of labile Cu^+^, it is plausible that the ion is unavailable for specific cellular processes within differentiating cells lacking *mCrip2*, which may also cause toxicity and impair cell differentiation.

### mCrip2 associates indirectly with chromatin in proliferating myoblasts, with interactions decreasing upon Cu supplementation and induction of differentiation

Considering the nuclear distribution patterns of mCrip2 and responsiveness to Cu supplementation in myoblasts, as well as the need of this protein for differentiation, we hypothesized that the protein may function as a transcriptional regulator within the nucleus. To investigate this, CUT&RUN was performed to determine the chromatin localization for mCrip2 under various conditions, including proliferation (48 h), differentiation (24 h), and distinct insulin- and Cu-supplementation conditions. Genome-wide analyses revealed that mCrip2 is more enriched on chromatin in proliferating cells relative to differentiating cells (**Fig. 7A,B**). We performed peak calling for mCrip2 in both proliferating and differentiating myoblasts using Sparse Enrichment Analysis for CUT&RUN (SEACR (113)). A total of 728 and 650 mCrip2 peaks were detected above IgG background in proliferating cells grown with or without Cu, respectively, with 274 peaks shared between these two conditions (**Fig. 7C**). As determined qualitatively by heatmap visualization, less peaks were identified in any of the three conditions differentiating myoblasts were culture. A total of 367 peaks were detected in differentiating myoblasts without insulin and Cu supplements (-Ins,-Cu), 389 peaks were identified in insulin-differentiated cells exhibited (+Ins), and 462 peaks were detected in Cu-supplemented differentiating myoblasts (+Cu; **Fig. 7D**).

**Figure 7.**
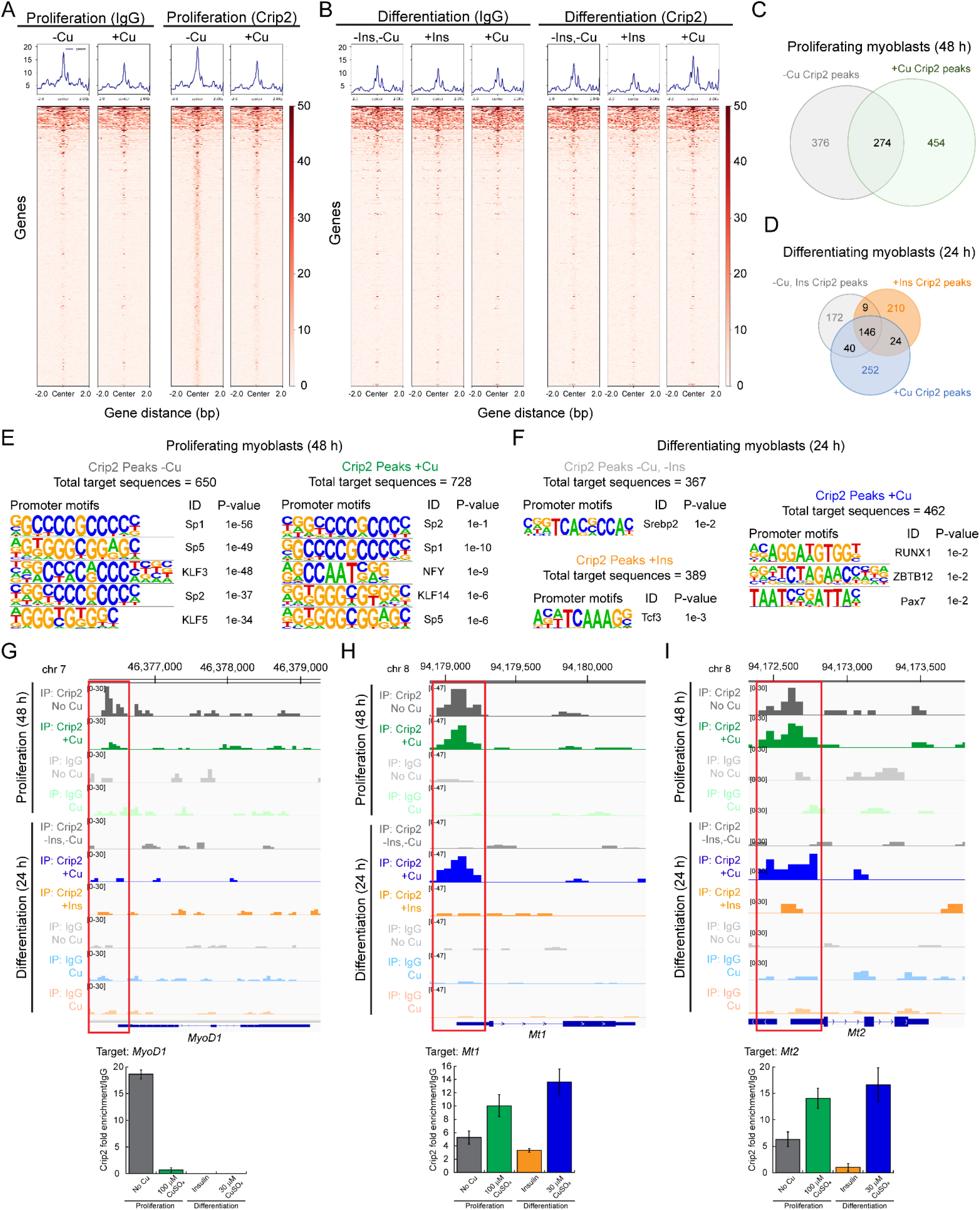
mCrip2 binding to chromatin in proliferating and differentiating myoblasts. Representative heat maps from peak calling by Sparse Enrichment Analysis (SEACR) for CUT&RUN. IgG controls and mCrip2 binding are depicted. (**A**) Proliferating WT primary myoblasts were supplemented or not with 100 µM CuSO_4_. (**B**) Differentiating (24 h) WT primary myoblasts were supplemented or not with insulin of 30 µM CuSO_4_. Overlap of CUT&RUN peaks of mCrip2 across the genome observed in proliferating **(C**) and differentiating **(D**) cells in the presence or absence of Cu and insulin. See complete set of genes in **Supp. Table 4**. Main changes of mCrip2 motif-binding dependent on Cu supplementation in proliferating and differentiating primary myoblasts. Novel consensus DNA-binding motifs identified from peak calling by Sparse Enrichment Analysis (SEACR) for CUT&RUN within mCrip2 peaks in proliferating cells **(E**) and differentiating myoblasts **(F**) supplemented or not with Cu. The top five most significant motifs enriched, including the DNA logo, its corresponding TF, and its P value are shown. See **Supp. table 4** for complete list of peaks. Representative genome browser tracks of CUT&RUN experiments examining mCrip2 binding to the *MyoD1* **(G**), *Mt1* **(H**) and *Mt2* **(I**) promoters in proliferating and differentiating myoblasts (upper panels). *MyoD1, M1 and Mt2* promoters were selected as a representative locus for validation by ChIP-qPCR (lower panels). Plots represent data obtained from 5 independent biological experiments.

To determine the underlying sequences for these peaks, we performed a *de novo* motif search on CUT&RUN peaks called for mCrip2 from proliferating and differentiating myoblasts cultured with or without Cu and insulin (**Fig. 7E,F**), focusing on the promoter-specific peaks. The majority of motifs identified correspond to TF-binding sites that contribute to muscle development and differentiation and interact with the classic myogenic markers *MyoD1* and *Myogenin*. Proliferation-specific motifs associated with mCrip2 localization were characterized by regions displaying high GC contents, with the top 5 motifs displayed (**Fig. 7E; Supp. Table 4**). In proliferating cells grown with or without Cu, SP1, SP2, and SP5 motifs were highly enriched, aligning with the high GC content, in addition to Krüppel-like factor (KLF) families of TF, which are implicated in regulating general metabolic and cellular fate pathways. Notably, the NFY (CCAAT-binding factor CBF) motif was only enriched in proliferating cells grown in the presence of Cu, supporting its role in transcription initiation control and chromatin modifier recruitment (147, 148). Fewer TF consensus motifs were detected for mCrip2 promoter-binding in differentiating myoblasts (**Fig. 7F**). In cells differentiated without insulin and Cu supplements (-Ins,-Cu), the only significantly enriched motif was for Srebp2, a TF known for its role in regulating lipid metabolism and energy homeostasis in liver, which has been proposed to also mediate these processes in muscle. For insulin-differentiated cells, the only significantly enriched motif was for Transcription Factor 3 (Tcf3), which has been shown to physically interact with MyoD1 and Myogenin, and to modulate their transcriptional activity (149). RUNX1, ZBTB12, and Pax7 motifs were detected in differentiating myoblasts supplemented with Cu. RUNX1 and ZBTB12 are expressed in myoblasts and are proposed to regulate the expression of genes necessary for muscle differentiation and regeneration by interacting with MyoD1 and Myogenin (150).

A gene-based analysis of mCrip2 binding locations identified several relevant peaks at genes related to transcriptional regulation, such as bromodomain proteins and Zn finger-containing TFs (**Supp. Table 4**). Additional examples include genes specific to the skeletal muscle lineage, genes related to maintenance of metal homeostasis, and genes involved in the assembly of vesicles/endosomes in both proliferating and differentiating cells. Specific peaks in proliferating myoblasts supplemented or not with Cu included chromatin regulators like the bromodomain and PHD finger containing 1 (*Brpf1*) and the Zn-binding protein *Zfp384*, which control histone binding and DNA transcription and *Kat6b*, an epigenetic regulator, which functions as a histone acetyltransferase.

Relevant to skeletal muscle development, *MyoD1,* an essential regulator of the skeletal muscle lineage, is also a target of mCrip2 in proliferating cells cultured in the absence of Cu (**Fig. 7G; Supp. Table 4**), hinting at the potential role of mCrip2 in muscle-related functions. Considering the relevance of MyoD1 in the skeletal muscle lineage, we confirmed the association of mCrip2 with the *MyoD1* promoter by ChIP-qPCR. Consistent with the CUT&RUN analyses, a robust interaction between mCrip2 and the *MyoD1* promoter in cells cultured in the absence of Cu was observed (**Fig. 7G**, lower panel), which decreased upon Cu supplementation in the medium, and upon induction of differentiation.

Additional targets involved in skeletal muscle development were identified. The regulator *Mef2D* (Myocyte Enhancer Factor 2D), emerged as a significant target of mCrip2 in actively dividing myoblasts exposed to Cu supplementation (**Supp. Table 4**). Other interesting peaks detected in genes related to skeletal muscle development targets of mCrip2 in proliferating cells were the proliferation marker *Pax7*, *DMPK* (myotonic dystrophy protein kinase), and the transcription factors *SP1* and *SP2.* SP1 and SP2 are known to regulate the expression of genes involved in muscle development and function, such as structural proteins, metabolic enzymes, and signaling molecules. It is noteworthy that binding motifs for these proteins were detected in proliferating myoblasts as well (**Fig. 7E**), suggesting a potential feedback loop between mCrip2 and the expression of these TFs. mCrip2 binding to structural genes was also detected in proliferating myoblasts, as exemplified by the *Mylpf* (myosin light chain, phosphorylatable, fast skeletal muscle or Mlc2) gene. Mylpf is an essential component of the myosin complex to enable muscle contraction and the proper functioning of fast-twitch muscle fibers.

A closer analysis of the CUT&RUN data revealed mCrip2 association to metal homeostasis-related genes in proliferating myoblasts and certain differentiation conditions (**Supp. Table 4**). For instance, mCrip2 targeted the Zn-exporter *Slc30a1* (*Znt1*) mainly in untreated proliferating cells. mCrip2 binding to 3’ UTR region of *Slc30a4* (*Znt4*) showed predominant association in both proliferating and differentiating cells. The promoter region of the Zn importer *Slc39a13* (*Zip13*) was also a mCrip2 target, specifically in proliferating cells. Genome browser track analyses showed mCrip2 enrichment over both *Mt1* and *Mt2* promoter regions in proliferating cells and in those differentiated with non-toxic levels of Cu. (**Fig. 7H,I**; upper panel highlighted in red; **Supp. Table 4**). These findings were corroborated through ChIP-qPCR analysis (**Fig. 7H,I**; lower panel). Notably, mCrip2 enrichment at the *Metallothioneins* promoters was approximately 2-fold higher in cells cultured with Cu compared to those without the metal. GO analyses for promoters showing mCrip2 occupancy and found diverse categories of genes enriched in each growth condition, highlighting possible diverse functional roles of mCrip2 in contributing to gene expression patterns during different stages of muscle cell development and under varying culture conditions (**Supp. Fig. 7; Supp. Table 4**).

### Transient interactions between mCrip2 and the chromatin remodeler Brg1

While mCrip2 is associated with a relatively small number of gene promoters, the functional diversity of its targets suggests that this protein plays a complex regulatory role in transcription across diverse cellular processes. The fact that mCrip2 belongs to the Class II family of LIM-domain proteins, which lack DNA-binding domains (151–155), coupled with the limited number of mCrip2 peaks across all tested culture conditions, led us to hypothesize an indirect functional role as transcriptional co-factor for this Cu-BP. As observed for other LIM-containing proteins, the Zn-finger regions may enable physical interactions with other proteins involved in transcription regulation. To test this hypothesis, we investigated the potential interaction between mCrip2 and Brg1, an essential chromatin remodeler in the skeletal muscle lineage (55, 56, 64, 156). Confocal microscopy analyses revealed partial colocalization of mCrip2 with Brg1 in proliferating primary myoblasts that increased in differentiating cells (**Supp. Fig. 8**). Unsuccessful immunoprecipitation attempts to detect the interaction between mCrip2 and Brg1 suggest a transient and weak association, which supports the hypothesis that mCrip2 acts as an indirect facilitator or activator for major transcriptional regulators.

### *mCrip2* deletion induced distinct transcriptional effects in proliferative and differentiating primary myoblasts

So far, CUT&RUN and confocal microscopy analyses suggest that mCrip2 is a potential facilitator or co-activator of transcription for essential genes, with *MyoD1* being a relevant target for myogenesis and *Metallothioneins* as important targets for metal response **(Fig. 7**). In addition, *mCrip2* KO impaired differentiation of primary myoblasts, likely due to the elevated levels of labile Cu observed (**Figs. 5,6**). These findings point to a role for mCrip2 in myogenesis and suggest an involvement in Cu handling within skeletal muscle cells. To gain deeper insights into the observed phenotypes of *mCrip2* KO in myogenesis at the transcriptional level, we conducted RNA-seq on myoblasts lacking *mCrip2* from proliferating (48 h) and differentiating (24 h) myoblasts. Principal component analyses (PCA) within each sample type showed that the replicates have minimal variability between replicates (**Supp. Fig. 9**) with differentially expressed genes included in **Supp. Table 5**.

In proliferating cells, *mCrip2* KO led to 1288 differentially expressed genes (DEGs) compared to control cells, with 624 genes upregulated and 664 downregulated. In differentiating myoblasts, a total of 4270 DEGs were detected, with 2523 genes upregulated, and 1747 genes downregulated. GO term analyses revealed that in proliferating cells, deletion of *mCrip2* led to decreased expression of genes related to mRNA processing, protein translation, ribosome synthesis, and chromatin organization and increased expression of genes related to diverse signaling pathways, cytoskeleton organization, and cell trafficking (**Fig. 8A**). We validated expression changes of representative genes using qRT-PCR. Specifically, we examined two genes related to ribosomal synthesis: RpsA and Rps8. RpsA is vital for 40S ribosomal subunit assembly and stability, as well as 20S rRNA precursor processing into mature 18S rRNA; Rps8 encodes a 40S subunit component (157, 158). Reduced mRNA levels in *mCrip2*-depleted myoblasts were consistent with RNA-seq analyses for both genes (**Fig. 8B**). RNA-seq and qPCR analyses in proliferating cells detected no changes in the expression of the *Pax7* gene (**Fig. 8B**), which agrees with the immunohistochemistry analyses presented above (**Fig. 4**).

**Figure 8.**
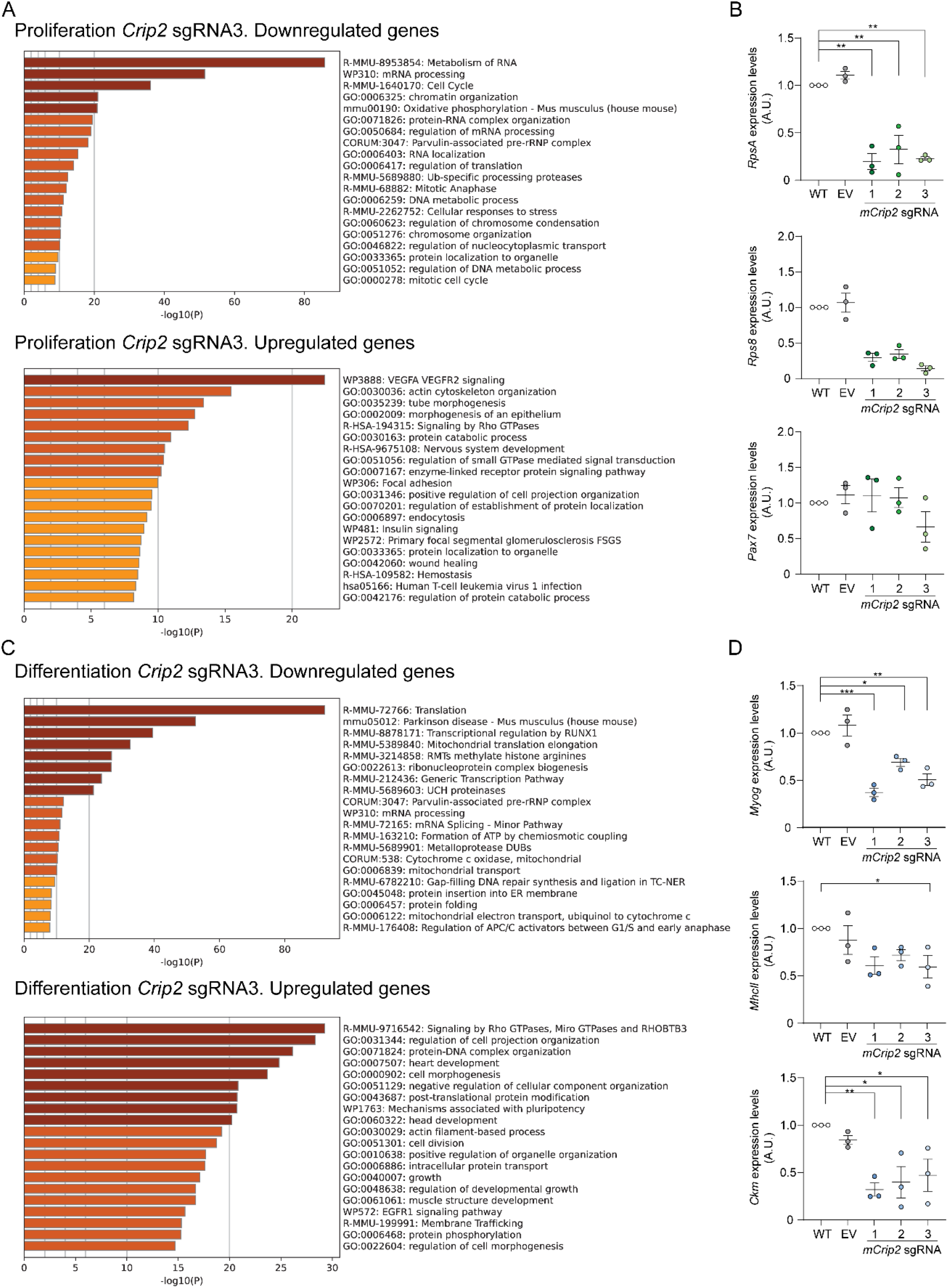
Changes in gene expression dependent on *mCrip2* knockout. GO term analysis of differentially expressed, down- and up-regulated genes induced by the KO of *mCrip2* (sgRNA3) and steady state mRNA expression of selected genes validated by qPCR in proliferating **(A,B**) and differentiating **(C,D**) myoblasts. Cut-off was set at 2.0 of the -log(adjusted P value). See **Supp. Table 5** for the complete list of genes. N = 3 All values are reported as means ± SEM. Significance was determined by two-way ANOVA; *p < 0.05, **p < 0.01, ***p < 0.001 and ****p <0.0001 compared to control cells.

In differentiating *mCrip2* KO myoblasts, reduced gene expression primarily affected processes linked to transcriptional regulation, mRNA processing, protein synthesis, and folding (**Fig. 8C**). A significant portion of downregulated genes were responsive to RUNX1 regulation, consistent with our motif search using CUT&RUN analyses for mCrip2 (**Fig. 7**). Additionally, *mCrip2* KO negatively impacted essential mitochondrial Cu-dependent events and mitochondrial transport-related genes during differentiation. Upregulated processes included signaling pathways like GTPase Rho, protein-DNA organization, cell division, growth, and mechanisms associated with pluripotency. Genes related to heart, head, and muscle structure development were also identified, with muscle structure development being the least significant among these three cellular types (**Fig. 8C**). Specifically in differentiating myoblasts, classic differentiation markers *Myogenin* (*Myog*), *myosin heavy chain II* (*MhcII*), and *muscle-specific creatine kinase* (*Ckm*) were downregulated in *mCrip2* KO cells, contributing to the observed differentiation defect at the transcriptional level (**Fig. 8D**). Focusing on metal-related genes, particularly metallothioneins *Mt1* and *Mt2*, RNA-seq analyses revealed significant reductions in *Mt1* expression and slight, insignificant decreases in *Mt2* expression in differentiating cells lacking mCrip2 (**Supp. Table 5**). Minor, non-significant reductions in *Mt1* expression were observed in proliferating myoblasts lacking *mCrip2* (**Supp. Table 5**). Notably, expression of *Mtf1*, the transcription factor responsible for *Mt* induction, remained unchanged upon *mCrip2* deletion, explaining the modest changes in metallothionein expression and supporting an indirect role for mCrip2 as a transcription facilitator. These findings reveal the impact of mCrip2 on important ribosomal, myogenic, and metalloprotective genes, highlighting its potential role in various molecular events governing muscle biology, while also suggesting a function as a Cu-responsive or Cu-regulating protein.

Using the RNAseq data, we performed a comprehensive approach to analyze transcriptional regulatory networks by integrating data from Transcriptional Regulatory Relationships Unraveled by Sentence-based Text-mining (TRRUST) with MetaScape (159). TRRUST provides curated transcriptional regulatory interactions sourced from literature, offering insights into the relationships between TFs and their target genes. These analyses revealed that the downregulated genes in proliferating *mCrip2* KO cells responded to TFs including Myc and Rbl1, while SP1 was the top TF identified for upregulated genes, a motif that was also identified in our CUT&RUN analyses (**Supp. Fig. 10A**). In differentiating KO myoblasts, genes regulated by Myogenin were among the most downregulated genes, followed by targets of Bcl3 and Twist2 (**Supp. Fig. 10B**). Among the genes that were induced, we identified Chd7, Trp53 and Sox4 as the potential TFs related to their expression.

To comprehensively dissect the molecular landscape of gene regulation by mCrip2 in proliferating and differentiating primary myoblasts, we integrated the CUT&RUN and RNA-seq datasets, to identify which DEGs were bound by mCrip2. With these integrated data, we found that in proliferating myoblasts cultured in the absence or presence of 73 and 64 genes were both DEGs in *mCrip2* KO myoblasts and were bound by mCrip2, respectively (**Supp. Table 6**). In differentiating cells, we detected few targets of mCrip2 that were DEG in differentiating myoblasts; only 2 genes were identified in cells cultured in the absence of insulin and Cu, 5 DEGs in cells differentiated with insulin and 14 DEG in cells supplemented with Cu (**Supp. Table 6**).

We examined whether these bound DEGs were enriched for any TF motifs (**Supp. Fig. 11**). Rfx5 (Regulatory Factor X 5), NRF (Nuclear Respiratory Factor), Hif1α (Hypoxia Inducible Factor 1 Alpha), ZNF652 (Zinc Finger Protein 652) and Tcf4 (Transcription Factor 4) emerged as the most significant promoter motifs in downregulated genes, particularly in cells supplemented with Cu. RFX5 is a regulator of the *MHC class II* gene that has been implied in muscle-specific gene expression. NRF governs mitochondrial protein-encoding genes, crucial for muscle function, while Hif1α responds to hypoxic stress in muscle tissue. Information regarding Znf652 in muscle is limited, yet its role as a Zn-binding protein suggests involvement in the mCrip2 regulatory network. Tcf4 is implicated in balancing myoblast proliferation and differentiation, along with maintaining muscle homeostasis and aiding in regeneration by modulating gene expression related to muscle growth and repair. Among the bound, upregulated DEGs in proliferating cells, NYF (Nuclear Factor Y) and ZNF322 (Zinc Finger Protein 322) motifs appeared in both proliferating myoblasts grown with or without Cu (**Supp. Fig. 11A,B**). NFY is proposed to regulate muscle-specific genes, while the role of ZNF322 in muscle has not been elucidated yet. KLF3, KLF14 and SP5 sites were detected in myoblasts cultured in the absence of Cu, while ZNF711, SP2, and ZNF652 were enriched in cells supplemented with Cu (**Supp. Fig. 11A,B**). As discussed above, KLF3, KLF14, SP5 and SP2 may regulate myogenesis, muscle fiber type specification, and metabolic gene expression, impacting muscle development and function. ZNF711 (Zinc Finger Protein 322) is another gene encoding a Zn finger protein, which similarly to other Zn fingers detected here has no specific function identified in muscle biology yet.

In myoblasts differentiated with insulin, the identified motifs for upregulated genes were CRE (cAMP Response Element) and KLF3 (**Supp. Fig. 11C**). CRE is a binding site for TFs like CREB, potentially promoting the expression of genes crucial for muscle growth and adaptation to exercise. For cells supplemented with Cu, LRF (Leukemia/lymphoma-related factor) was associated with downregulated genes, while MYB (Myeloblastosis Oncogene) and ERRα (Estrogen-Related Receptor Alpha) were linked to upregulated genes (**Supp. Fig. 11D**). Although the roles of LRF and MYB in skeletal muscle differentiation are not fully understood, they may impact muscle development through their influence on cell cycle regulation, differentiation, and apoptosis. Additionally, ERRα, a nuclear receptor TF, regulates energy metabolism and mitochondrial biogenesis, potentially enhancing endurance and fatigue resistance by promoting oxidative metabolism in muscle fibers.

### mCrip2 is a Cu-responsive transcriptional modulator of metalloprotective genes

Considering the elevated expression and interaction of mCrip2 with *Mt1* and *Mt2* promoters upon Cu supplementation, alongside the observed accumulation of Cu in myoblasts lacking the gene, we propose that mCrip2 functions as a metal-responsive protein that facilitates a response to excess metal accumulation. This hypothesis is supported by the potential mechanism wherein mCrip2 deficiency leads to impaired expression of metallothioneins upon metal supplementation. To test this, we examined Mt1 expression and localization in proliferating and differentiating primary myoblasts cultured with or without Cu (**Fig. 9**). Western blot analyses revealed induction of Mt1 in both proliferating and differentiating myoblasts upon exposure to non-toxic Cu levels (**Fig. 9A,B**). Notably, deletion of *mCrip2* prevented the increase in Mt1 expression in Cu-cultured cells. Consistent with this, confocal microscopy demonstrated elevated Mt1 levels in control cells under non-toxic Cu conditions, while *mCrip2* KO cells failed to exhibit such induction (**Fig. 9C,D**). Together these results demonstrate that mCrip2 may act as a transcriptional regulator of essential metalloprotective genes upon exposure to metals.

**Figure 9.**
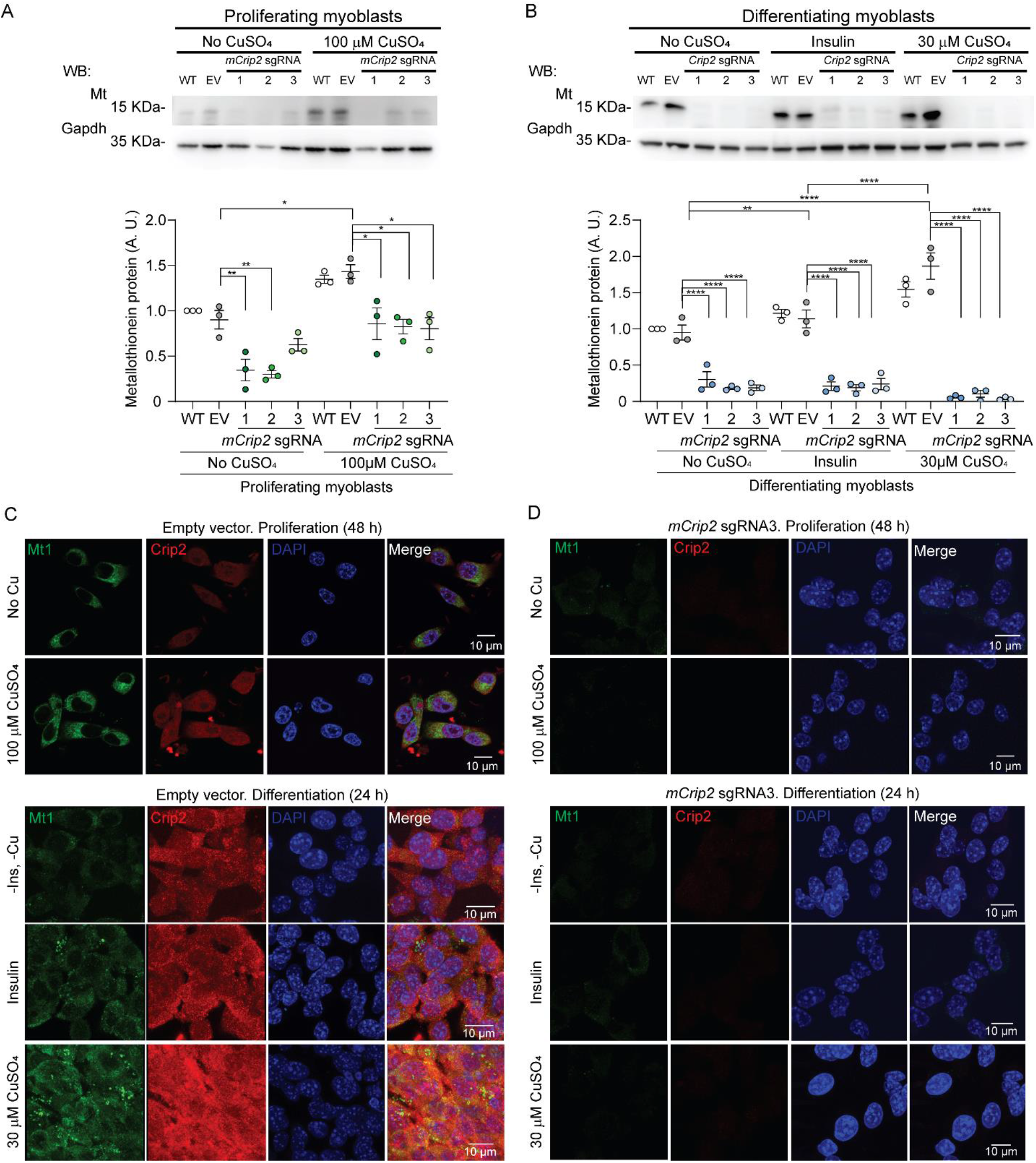
Depletion of *mCrip2* prevents the Cu-dependent induction of metallothioneins. Representative western blot (upper panel) and quantification (bottom panel) of Metallothionein 1 (Mt) expression in control and *mCrip2* KO proliferating (48 h; **A**) and differentiating (24 h; **B**) primary myoblasts. Gapdh was used as loading control. Representative confocal microscopy of Metallothionein 1 (Mt1, green) and mCrip2 (red) expression of empty vector (**C**) and the *mCrip2* sgRNA3 myoblast clone (**D**) cultured under the same conditions as in A. Nuclei were stained with DAPI (blue). The images depicted are representative of 3 independent biological experiments. Size bar = 10 µm.

## DISCUSSION

### mCrip2 is a Cu-responsive regulator in skeletal muscle cells

In this work we characterized the contributions of the Cu^+^-binding, LIM-domain protein mCrip2, in the development of primary myoblasts derived from mouse satellite cells. Within the landscape of cell determination and differentiation, the LIM motif has emerged as an essential unique double-Zn-finger structure required for the function of several proteins for these developmental processes (151, 160). Consistent with previous research, our study demonstrates that mCrip2 is a high-affinity Cu^+^-binding protein (5). Here, we showed that in primary cultures of skeletal muscle cells Cu supplementation enhances mCrip2 expression. In contrast, investigations on H1299 lung cancer cells reveal decreased levels of both endogenous and recombinant hsCRIP2 upon Cu exposure, countered by the chelator BCS (5). *In vitro* thermal stability assays pointed to a Cu^+^-mediated reduction in hsCRIP2 stability, corroborated by degradation assays with CHX suggesting ubiquitin/proteasome-mediated degradation of Cu-bound hsCRIP2, and an increase in autophagic activity upon *hsCRIP2* KD(5). Our differentiation model of primary myoblasts revealed a different regulation of mCrip2 dependence on Cu, in both nuclear and cytosolic fractions. Our studies further pointed to potential lineage-specific roles in transcriptional regulation and metal sensing for mCrip2. In the murine myoblast nuclei, mCrip2 seems to have an indirect regulatory role of transcription, potentially by facilitating the recruitment of elements of the transcriptional machinery, such as chromatin remodelers, to chromatin in a Cu-dependent manner. These findings point to Cu-dependent lineage-specific functions of mCrip2, illuminating novel regulatory mechanisms governed by Cu that render differential functions specific to cell lineages, shedding light on alternative potential regulatory mechanisms influenced by Cu-BP in cellular homeostasis.

### The regulatory role of Crip2 in the nuclei of primary myoblasts derived from mouse satellite cells

Numerous LIM proteins exert their influence on developmental events by modulating gene expression (161–163, 170–173). LIM proteins lacking a DNA-binding homeodomain, such as Crip2, are thought to play a role in the transcriptional regulation during cell differentiation. Proto-oncogenes RBTN1 and RBTN2, activated in T-cell leukemia, exemplify this phenomenon. Mice with null mutations in the *Rbtn2* gene die early in development due to the absence of erythroid precursors, highlighting the essential role of RBTN2 in normal erythroid development (153). TAL1, a key bHLH transcription factor in erythroid development, co-expresses and interacts with RBTN2 and RBTN1 through their respective LIM and bHLH domains in erythroid cells (172, 173). Similar synergistic interactions have been observed in pancreatic cells, where the LIM protein lmx-1 and the bHLH protein E47 interact to activate the expression of the insulin gene (171). Crip3 illustrates yet another instance of these cooperative interactions, engaging in nuclear protein interactions with muscle-specific bHLH TFs, including MyoD1, MRF4, and myogenin (170).

In line with these observations, our CUT&RUN analyses showed that mCrip2 interacts with a selected group of genes with highly specific and essential roles in myogenesis, transcriptional regulation, metal homeostasis, and endosome formation. RNA-seq analyses further demonstrated the broader impact of potentially direct and indirect interactions of mCrip2 with either specific promoters or with major transcriptional regulators. Our data agrees with the hypothesis that the interactions between tissue-specific bHLH factors and LIM proteins may signify a common mechanism employed by diverse developmental systems to enhance transcriptional activity. Supporting a role as a transcriptional facilitator during skeletal muscle development, we found mCrip2 binds to promoter regions of few, but relevant genes associated to myogenesis, such as *MyoD1*. Another interesting gene promoter was *Brpf1*, which like *MyoD1* was a target of mCrip2 in proliferating cells, with an interaction decreasing upon induction of differentiation. Brpf1 is a multivalent chromatin regulator possessing three histone-binding domains, one non-specific DNA-binding module, and several motifs for interacting with and activating three lysine acetyltransferases. Brpf1 mutations have been linked to intellectual disability syndromes, where important muscle defects, such as hypotonia, have been observed in patients (174). The promoter of *Mef2D*, or Myocyte Enhancer Factor 2D, was another important TF target of mCrip2 in proliferating myoblasts supplemented with Cu. Mef2D is expressed in skeletal, cardiac, and smooth muscle tissues, highlighting its role in the regulation of gene expression in various muscle types (175). Mef2D levels rise during myogenic differentiation, driving muscle-specific gene expression, and it is involved in the determination of muscle fiber type (176). It regulates the expression of genes associated with specific muscle fiber types, contributing to the diversity of muscle fibers with distinct contractile and metabolic properties. Mef2D interacts with other transcription factors, such as MyoD1 and Myogenin, forming a regulatory network that orchestrates the complex myogenic process. Dysregulation of Mef2D expression or activity is associated with various muscle-related disorders, including muscular dystrophies and cardiomyopathies.

Another important observation supporting the function of mCrip2 as a facilitator of transcription and Cu-responsive function, is the binding to promoters of the metallothionein genes, *Mt1* and *Mt2*. The fact that mCrip2 was found at the promoters of these genes in proliferating cells cultured with basal media, and in cells where Cu was supplemented, highlights the importance of mCrip2 in facilitating transcription of homeostatic genes, and a role as a potential metal sensing protein as well. Experiments have addressed the role of *Mt* genes in myogenesis. Early studies showed upregulation of *Mt* expression in aging skeletal muscles in both rats and humans (177, 178). Additionally, upregulation of Mt genes has been shown in various types of skeletal muscle atrophy (179, 180). Notably, in *Sod1* KO mice undergoing aging and denervation, the levels of *Mt1* and *Mt2* are elevated in the gastrocnemius muscle, indicating a potential protective function against oxidative stress in the absence of the dismutase (181). Conversely, deficiency in *Mt1* and *Mt2* genes led to increased myogenesis in *Mt* KO mice (182). In fact, a recent study explored the role of ubiquitously expressing Mt on muscle differentiation. CRISPR-Cas9 mediated deletion of *Mt1* and *Mt2* genes in C2C12 myoblasts showed that deficiency of these genes promotes myocyte fusion, with *MyoD1* and *myogenin* upregulated at the late stage of myotube differentiation (183). Furthermore, while the expression of fast-twitch MHC protein expression remained stable, the expression of slow-MyHC expression was higher in KO cells than controls (183). This effect was reverted by the antioxidant N-acetylcysteine, suggesting that the antioxidant effects of Mts may be involved in slow-twitch myofiber formation in skeletal muscle (183). Although an important function for mCrip2 as a nuclear protein has emerged as a transcription co-adaptor protein (184), further research into the nuclear function of mCrip2 in response to Cu-supplementation during skeletal muscle differentiation is needed to fully understand the multifaceted role of this factor during myogenesis.

We propose that mCrip2 likely assists in recruiting additional components of the transcriptional machinery, such as the SWI/SNF complex, to activate specific myogenic genes like *MyoD1* or *Mef2D* in a Cu-dependent manner. Previous studies from our group have shown that Cu plays multiple regulatory roles during skeletal muscle differentiation, promoting the expression of myogenic genes (9, 17, 41). Therefore, it is possible that Cu supplementation may activate alternative transcriptional networks where mCrip2 binds specifically to myogenic promoters in proliferating cells. As differentiation begins and other essential factors bind to the promoters, the transcriptional role of mCrip2 may be fulfilled, leading to its detachment from those specific loci. The hypothesis of mCrip2 acting as an indirect facilitator of transcription responsive to Cu is also supported by the observation that mCrip2 is recruited to the promoter region of metallothionein genes upon metal supplementation. Thus, we propose that mCrip2 may act as a Cu-responsive protein regulating the expression of crucial responders to various stimuli, including differentiation cues and metal handling.

### Disruption of Cu homeostasis in *mCrip2*-deficient cells results in muscle differentiation defects

Excessive Cu levels can profoundly impact muscle differentiation by disrupting key cellular processes essential for myogenesis (17). Firstly, Cu-induced oxidative stress can damage critical cellular components such as proteins, lipids, and DNA, which are necessary for proper muscle cell differentiation. Oxidative damage can impair signaling pathways and transcriptional regulation networks involved in controlling the expression of myogenic genes, thereby disrupting the progression of muscle differentiation programs. Moreover, Cu overload can dysregulate the fine balance of additional metals, such as Zn and Fe, within muscle cells. These disruptions can interfere with the activity of metal-dependent enzymes and transcription factors crucial for muscle development and differentiation. For example, Zn-finger TFs play pivotal roles in regulating the expression of muscle-specific genes during myogenesis, and alterations in Zn availability due to Cu overload can impair their function. Furthermore, excessive Cu can perturb the delicate redox balance present in developing muscle tissue, leading to aberrant activation or inhibition of redox-sensitive signaling pathways involved in muscle differentiation. Redox signaling plays a fundamental role in modulating various aspects of myogenesis, including cell proliferation, differentiation, and survival. Disruption of these pathways by Cu-induced oxidative stress can compromise the coordination of molecular events necessary for muscle cell differentiation. Assessment of cytosolic and nuclear Cu levels by AAS revealed an excess of the ion in myoblasts depleted of *mCrip2*, which were corroborated by confocal microscopy analyses indicating changes in labile Cu^+/2+^ pools. Increased concentrations of labile Cu intensify oxidative stress due to its ability to generate ROS. These findings reveal the necessity for endogenous antioxidants and Cu chelators, like metallothioneins, in buffering intracellular metals and protecting against oxidative stress. The observed decrease in the labile Cu^+^ pool in wild type myoblasts undergoing differentiation suggests an increased sequestration of this metal. Conversely, *mCrip2*-deficient myoblasts had significantly elevated labile Cu^+^ and Cu^2+^, and additionally, Cu supplementation partially rescued myoblast detachment suggesting an enhanced usage of this metal as a compensatory mechanism for reduced metal regulation by mCrip2. Further studies are needed to elucidate how mCrip2 modulates Cu levels in skeletal muscle cells and to clarify its role as a metal sensing protein. In conclusion, mCrip2 emerges as an important and intricate regulator in skeletal muscle development and cellular homeostasis maintenance.

## AUTHOR CONTRIBUTIONS

Odette Verdejo-Torres: Data curation; Formal analysis; Investigation; Visualization; Validation; Writing—review and editing

David C. Klein: Data curation; Formal analysis; Investigation; Visualization; Writing—review and editing

Lorena Novoa-Aponte: Data curation; Formal analysis; Investigation; Visualization; Writing— review and editing

Jaime Carrazco-Carrillo: Investigation; Writing—review and editing

Denzel Bonilla-Pinto: Investigation; Validation; Writing—review and editing

Antonio Rivera: Investigation; Writing—review and editing

Fa’alataitaua Fitisemanu: Investigation; Writing—review and editing

Martha L. Jiménez-González: Investigation

Lyra Flinn: Investigation

Aidan T. Pezacki: Methodology; Writing—review and editing

Antonio Lanzirotti: Investigation; Methodology; Writing—review and editing

Luis Antonio Ortiz-Frade: Data curation; Formal analysis; Methodology; Writing—review and editing

Christopher J. Chang: Resources; Formal analysis; Methodology; Writing—review and editing; Funding acquisition

Juan G. Navea: Data curation; Formal analysis; Investigation; Methodology; Writing—review and editing; Funding acquisition; Supervision

Crysten Blaby-Haas: Formal analysis; Investigation; Visualization; Methodology; Writing—review and editing; Funding acquisition

Sarah J. Hainer: Data curation; Formal analysis; Investigation; Visualization; Methodology; Writing—review and editing; Funding acquisition; Supervision

Teresita Padilla-Benavides: Conceptualization; Data curation; Formal analysis; Investigation; Visualization; Methodology; Writing—original draft; Project administration; Funding acquisition; Supervision

## ACKNOWLEDGEMENTS

This work was supported by NIH grants NIAMS-R01AR077578 (to T.P.-B.), GM79465 (to C.J.C), and R35GM133732 (to S.J.H). C.J.C. is a CIFAR Fellow. J.G.N is supported by the Henry Dreyfus Teacher-Scholar Awards Program. This project used the Illumina NextSeq2000 available at the University of Pittsburgh Health Sciences Sequencing Core at UPMC Children’s Hospital of Pittsburgh for sequencing, with special thanks to its director, William MacDonald. This research was supported in part by the University of Pittsburgh Center for Research Computing, RRID:SCR_022735, through the resources provided. Specifically, this work used the HTC cluster, which is supported by NIH award number S10OD028483. Portions of this work were performed at GeoSoilEnviroCARS (The University of Chicago, Sector 13), Advanced Photon Source (APS), Argonne National Laboratory. GeoSoilEnviroCARS was supported by the National Science Foundation – Earth Sciences (EAR – 1634415). This research used resources of the Advanced Photon Source, a U.S. Department of Energy (DOE) Office of Science User Facility operated for the DOE Office of Science by Argonne National Laboratory under Contract No. DE-AC02-06CH11357. Work at the Molecular Foundry was supported by the Office of Science, Office of Basic Energy Sciences, of the U.S. Department of Energy under Contract No. DE-AC02-05CH11231 (CEB-H). The authors are thankful to Dr. Pablo Reyes-Gutierrez, Ms. Natalie O’Neill & Mr. Michael Quinteros for technical assistance. We thank Dr José Argüello’s laboratory at Worcester Polytechnic Institute for facilitating access to their Atomic Absorption Spectrometer and to Dr. Stephanie Weiner & Mr. Ahmed Almohamed for critical discussions of our work.

