## Supplementary information for "Cysteine Rich Intestinal Protein 2 is a copper-responsive regulator of skeletal muscle differentiation"

^2^ Department of Biological Sciences. University of Pittsburgh, Pittsburgh, PA. 15207. USA.

^3^ Present address: Genetics and Metabolism Section, Liver Diseases Branch, National Institutes of Diabetes and Digestive and Kidney Diseases, National Institutes of Health, Bethesda, MD. USA

^4^ Centro de Investigación y Desarrollo Tecnológico en Electroquímica, Querétaro. México

^5^ Chemistry Department. Skidmore College, Saratoga Springs New York, 12866. USA

^6^ Department of Chemistry. University of California, Berkeley, California, 94720. USA

^7^ Center for Advanced Radiation Sources, The University of Chicago, Lemont, IL 60439. USA

^8^ Department of Molecular and Cell Biology. University of California, Berkeley, California, 94720. USA

^9^ Molecular Foundry, Lawrence Berkeley National Laboratory, Berkeley, CA, USA & DOE Joint Genome Institute, Lawrence Berkeley National Laboratory, Berkeley, CA. USA

^&^ Present address: Progenra, Inc. Malvern PA 19355. USA

**Emails and ORCID:**

**List of supplementary materials**

**Supplementary figures**

Supplementary Figure 1. Phylogenetic analyses of the CRIP family members and appearance of the Cu+-binding sites.

Supplementary Figure 2. Maximum likelihood tree of individual LIM domains from the CRIP family.

Supplementary figure 3. Biochemical characterization of hsCRIP2.

Supplementary Figure 4. CRISPR/Cas9-mediated KO of Crip2 impairs differentiation of cultured primary myoblasts.

Supplementary Figure 5. CRISPR/Cas9-mediated KO of mCrip2 impairs differentiation of cultured primary myoblasts.

Supplementary Figure 6. Myoblasts depleted of mCrip2 present elevated levels of labile Cu+/2+ pools.

Supplementary Figure 7. GO term analyses of Crip2 annotated peaks in both proliferating and differentiating primary myoblasts

Supplementary Figure 8. mCrip2 partially co-localizes with Brg1 in primary myoblasts.

Supplementary Figure 9. Principal Component Analysis (PCA) plot showing the variance of the control and Crip2 sgRNA datasets used in the RNAseq analyses

Supplementary Figure 10. TRRUST gene set enrichment analysis

Supplementary ¬¬¬Figure 11. Main changes of mCrip2 motif-binding dependent on Cu supplementation in proliferating and differentiating primary myoblasts.

**Supplementary tables:**

Supp. Table 1. Crip2 protein sequence analyses.

Supp. Table 2. List of primers used in this study.

Supp. table 3. Parameters obtained from Cyclic voltammetry at different scan rates for Au bare and cysteine modified electrodes with hsCRIP2 protein.

Supp. Table 4. CUT&RUN mCrip2 Annotated peaks. All conditions

Supp. Table 5. RNAseq *mCrip2* KO DEG Prol and Diff.

Supp. Table 6. CUT&RUN – RNAseq merge analyses.

**SUPPLEMENTARY FIGURES**

**Supplementary Figure 1**
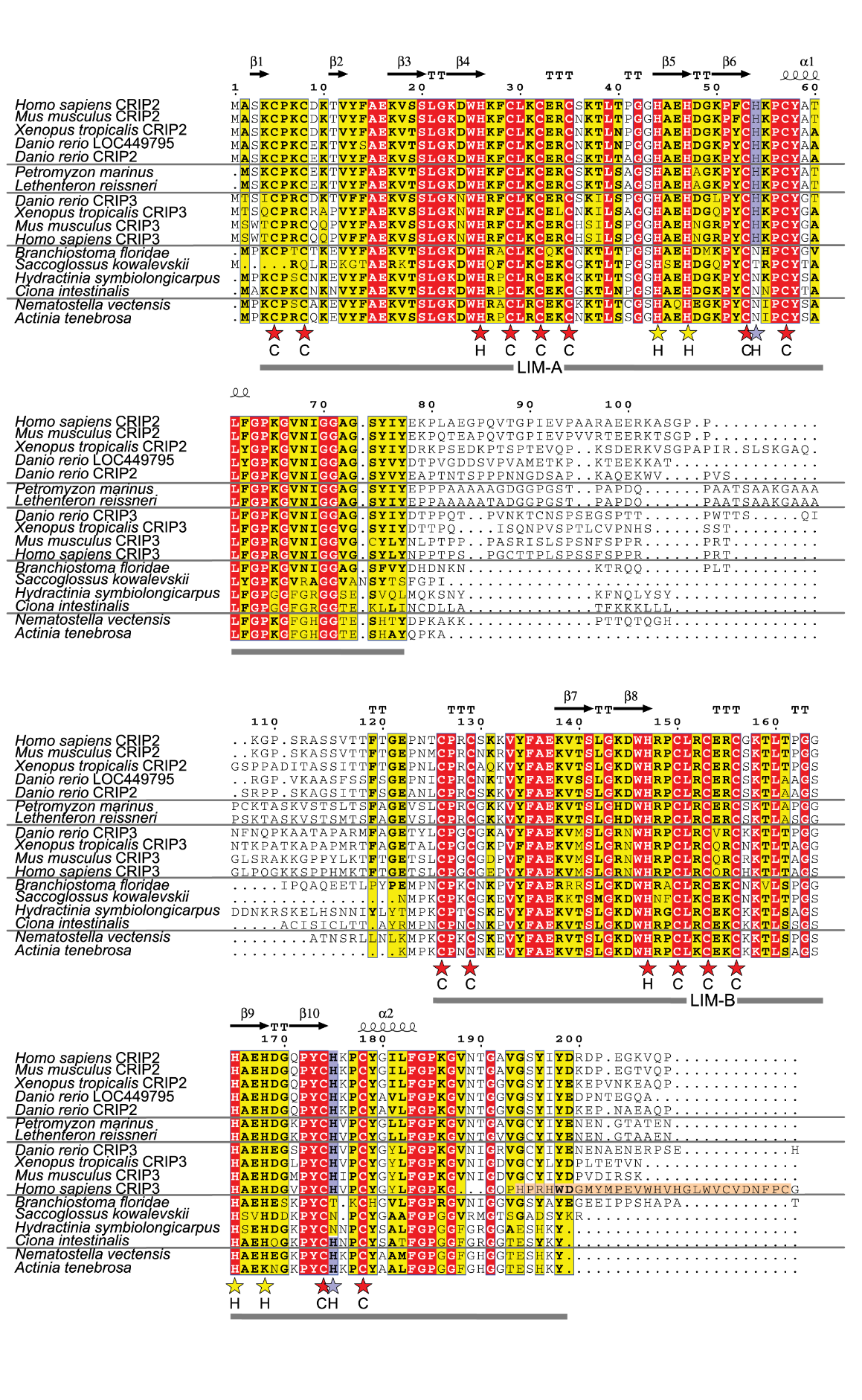

**Supplementary Figure 1. Phylogenetic analyses of the CRIP family members and appearance of the Cu^+^-binding sites.** Multiple sequence alignment of two-domain CRIPs. The Alpha Fold-predicted secondary structure of human CRIP2 is given at the top. Columns with white font and red shading are strictly conserved positions. Yellow shading represents a similarity score above 0.7. Purple shading and purple stars highlight the CRIP2/CRIP3-specific histidine residue near the Zn2 sites of LIM-A and LIM-B. Red stars are used to highlight the residues involved in Zn-binding. Yellow stars highlight the CRIP-specific histidine residues in the HxxH motif.

**Supplementary Figure 2**

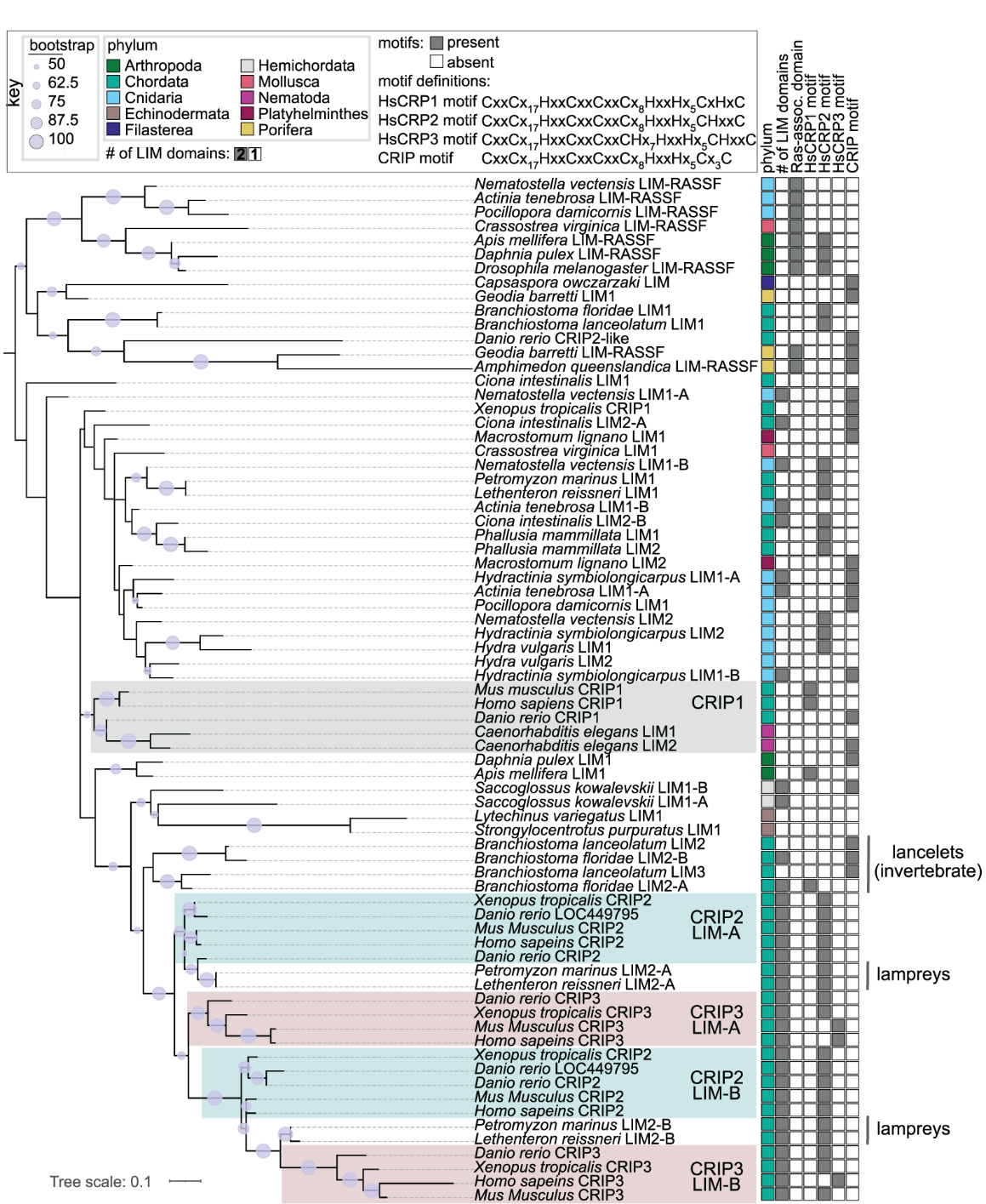

**Supplementary Figure 2. Maximum likelihood tree of individual LIM domains from the CRIP family.** For *Homo sapiens*, *Mus musculus*, *Xenopus tropicalis*, and *Danio rerio* the organism name is followed by the annotated protein name. For all other species, the CRIP homolog is labeled with “LIM” followed by a number. In all cases, if the LIM domain is from a two-LIM domain protein, the protein name is followed by “-A” to designate the N-terminal LIM domain or “-B” to designate the C-terminal LIM domain. The panel to the right is colored according to the key. The CRIP1 clade containing human and mouse CRIP1 is shaded in grey. The clades containing CRIP2 LIM domains are shaded teal and clades containing CRIP3 LIM domains are shaded peach.

**Supplementary Figure 3**

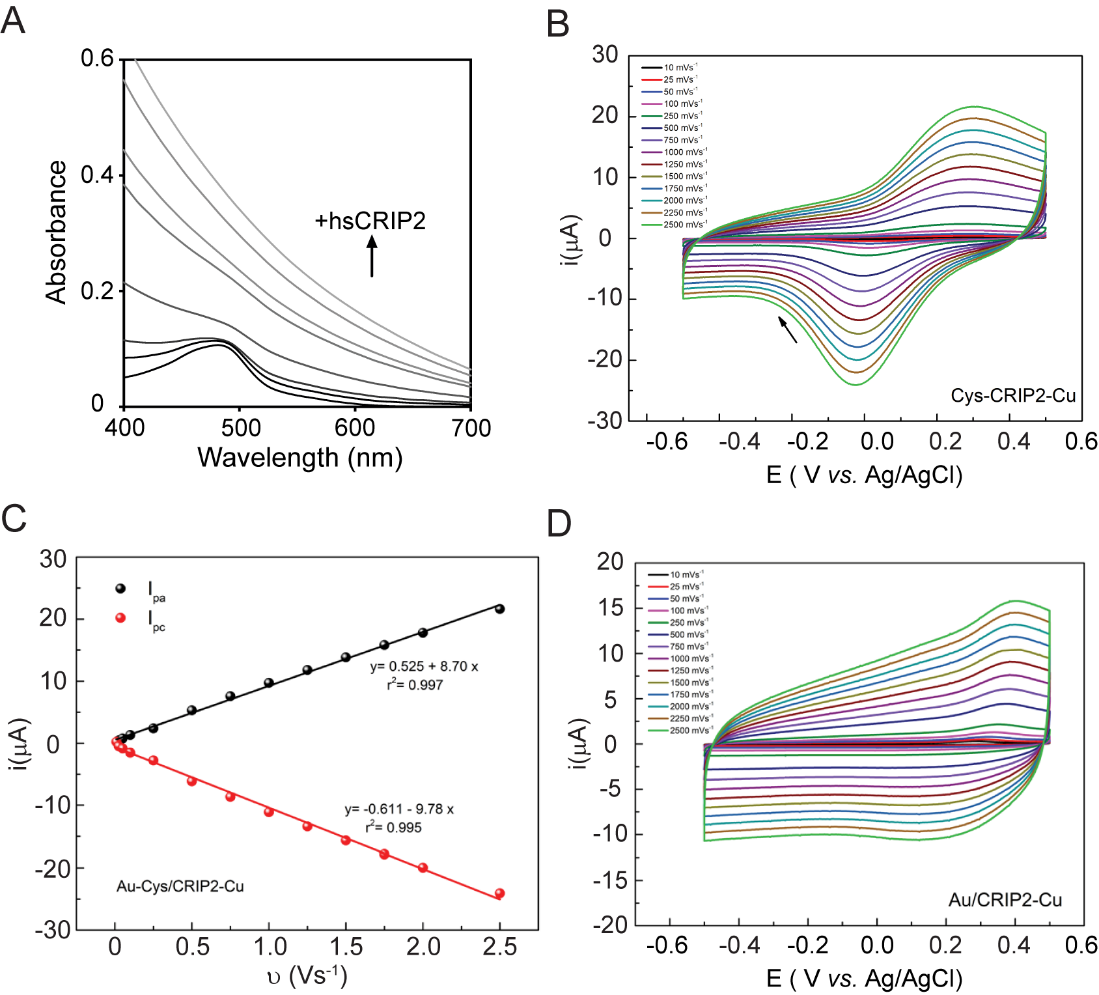

**Supplementary figure 3. Biochemical characterization of hsCRIP2. A.** Determination of the Cu^+^/Zn^2+^ dissociation constant (K_D_) of hsCRIP2 using the BCS competition assay. Spectrophotometric titration of 25 µM BCS and 10 μM Cu^+^ and Zn^2+^ with increasing concentrations of hsCRIP2. Protein precipitation was detected upon incubation with both metals impeding the changes in absorbance at 483 nm. **B.** Cyclic voltammograms obtained for the Au-Cys/hsCRIP2-Cu electrode in solution at different scanning rates (10 to 2500 mV s^-1^). **C.** Peak currents *vs*. scanning rate of the voltammograms obtained for Au-Cys/hsCRIP2-Cu. **D.** Cyclic voltammograms obtained for the Au/hsCRIP2-Cu electrode in solution at different scanning rates (10 to 2500 mV s^-1^).

**Supplementary Figure 4**

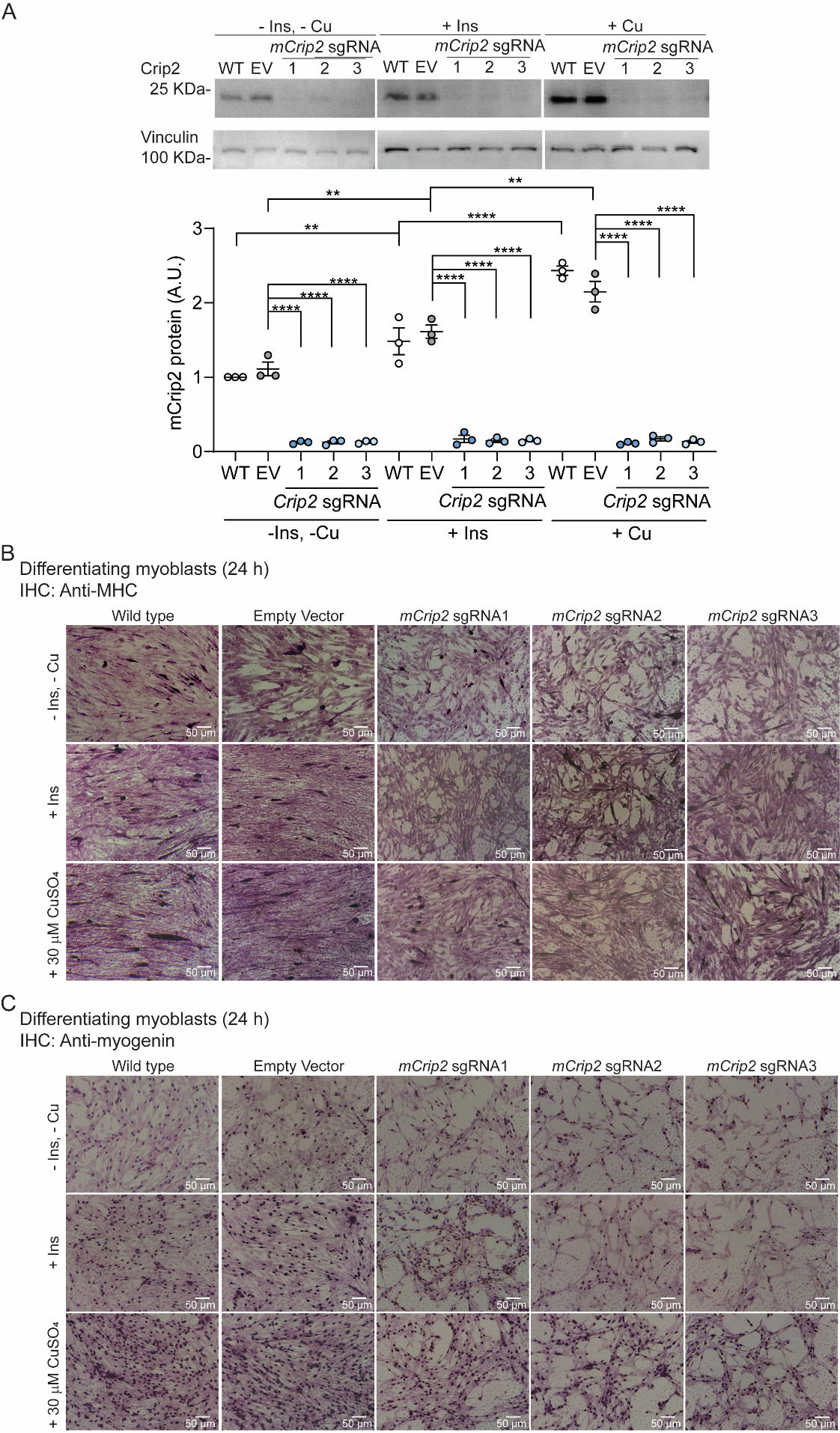

**Supplementary Figure 4. CRISPR/Cas9-mediated KO of *Crip2* impairs differentiation of cultured primary myoblasts. (A)** Representative western blot (upper panel) and quantification (lower panel) of Crip2 expression in wild type primary myoblasts and myoblasts transduced with either empty vector or three different sgRNAs against *Crip2* (1, 2, 3) obtained 24 h after inducing differentiation in the presence or absence of insulin, Cu, as indicated. Vinculin was used as a loading control. Representative light micrographs of differentiating myoblasts immunostained for myosin heavy chain **(B)** and myogenin **(C)** at 24 h in the presence or absence of insulin and Cu. Data represent three independent biologic experiments +/- SE. **P < 0.01; ****P < 0.0001.

**Supplementary Figure 5**

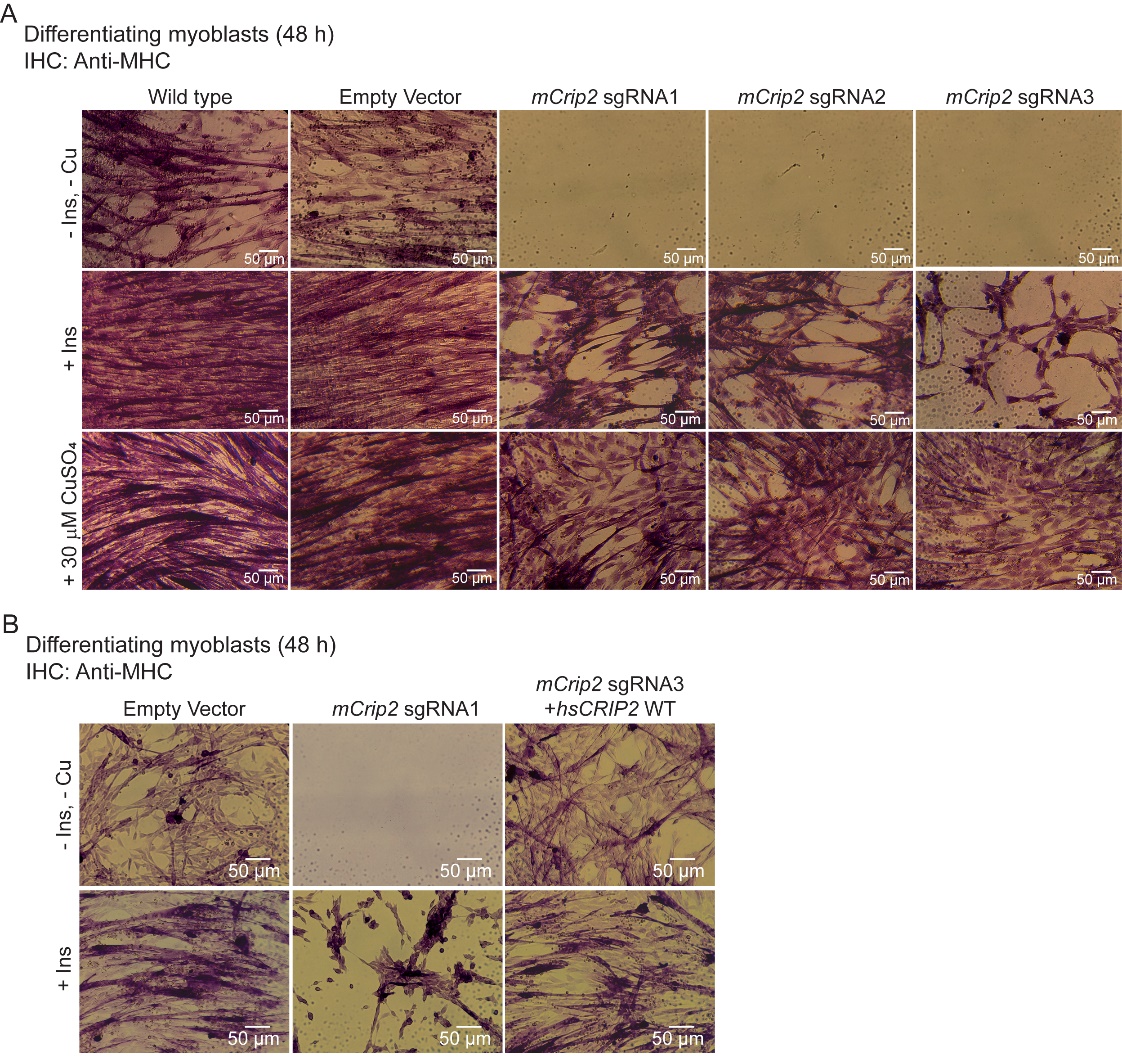

**Supplementary Figure 5.** **CRISPR/Cas9-mediated KO of *mCrip2* impairs differentiation of cultured primary myoblasts. (A)** Representative light micrographs of differentiating myoblasts immunostained for myosin heavy chain (MHC) after 48 h of inducing differentiation in the presence or absence of insulin and Cu. **(B)** The differentiation defect of *mCrip2* KO myoblasts is rescued by reintroducing *hsCRIP2* gene. Representative light micrographs of differentiating myoblasts immunostained for MHC. Images are represents 3 independent biological experiments.

**Supplementary Figure 6**

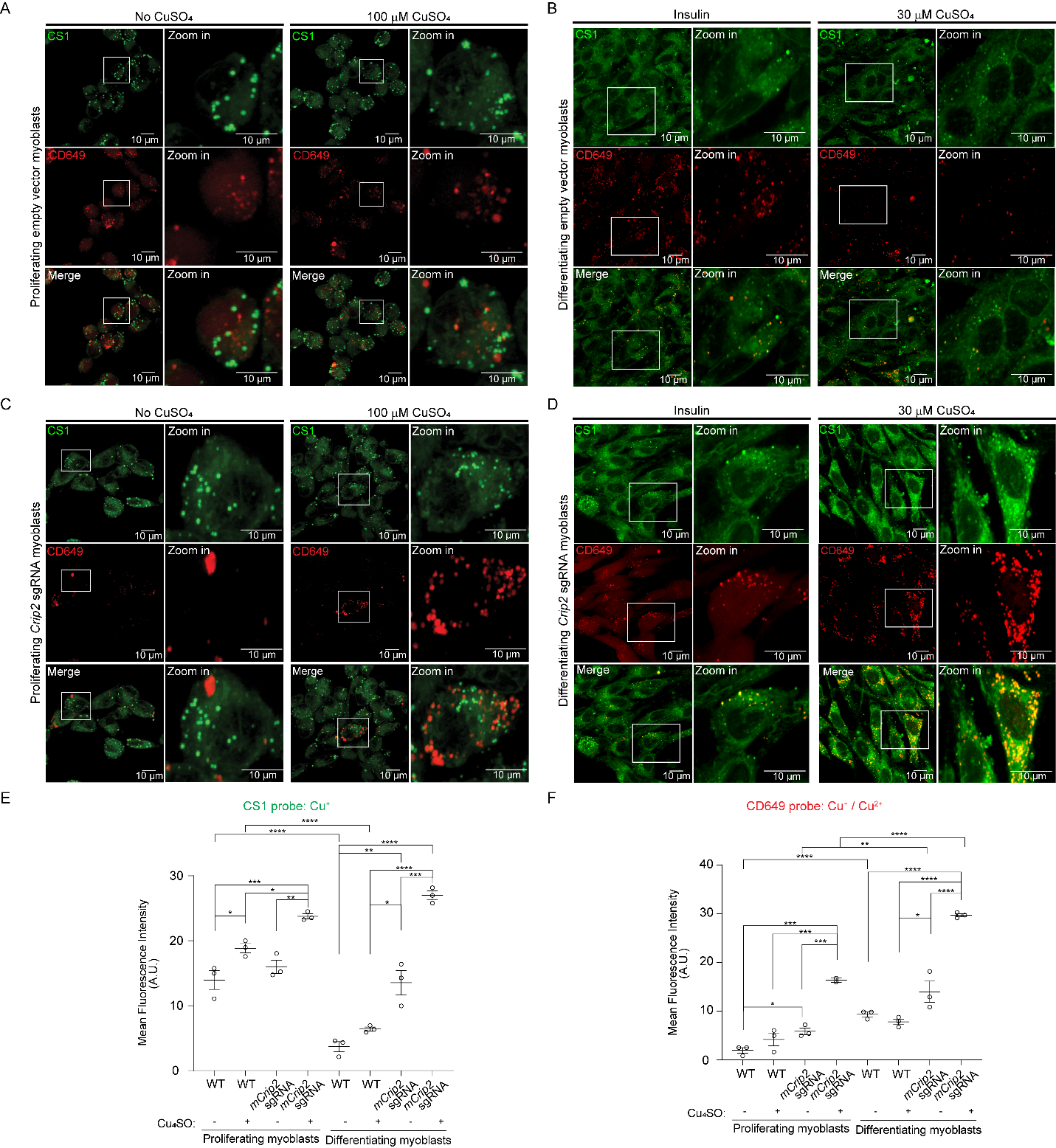

**Supplementary Figure 6. Myoblasts depleted of *mCrip2* present elevated levels of labile Cu^+/2+^ pools.** Live-cell confocal microscopy images of wild type primary myoblasts and KO for m*Crip2* supplemented or not with 100 µM CuSO_4_. Cu^+^ imaging was performed by incubating the cells with 5 µM CS1 (Cu^+^, green) and CD649 (Cu^+/2+^, red) for 10 min at 25 ºC and. Wild type proliferating **(A)** and differentiating myoblasts **(B)**. Primary myoblasts with CRISPR/Cas9-mediated deletion of m*Crip2* under proliferation **(C)** and differentiating conditions **(D)**. Quantification of the fluorescence of live-cell imaging for Cu^+^ with the CS1 probe (**E**) and Cu^+/2+^ with the CD649 probe **(F)** in proliferating myoblasts and differentiating myoblasts. N=3, *P < 0.05; **P < 0.01; ***P < 0.001; ****P < 0.0001.

**Supplementary Figure 7**

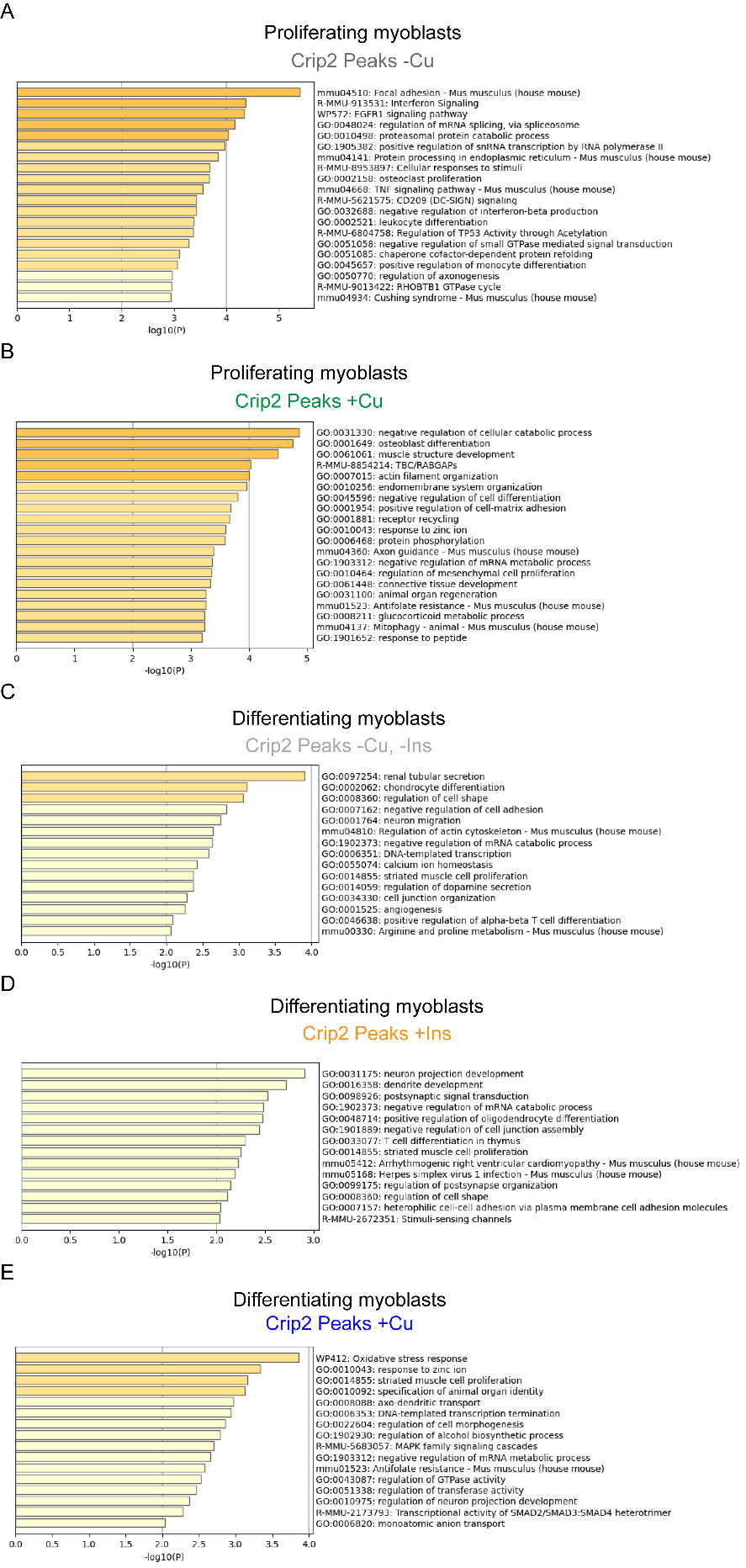

**Supplementary Figure 7. GO term analyses of Crip2 annotated peaks in both proliferating and differentiating primary myoblasts**. Heatmaps of enriched terms across Crip2-bound genes obtained with Metascape from peak calling by Sparse Enrichment Analysis (SEACR) for CUT&RUN. Proliferating cells grown in the absence of Cu (**A**) or supplemented with 30 µM CuSO_4_ (**B**). Myoblasts differentiated in the absence of Insulin and Cu (**C**) and supplemented with insulin (**D**) or with Cu (**E**). See **Supp. Tables 4** for the complete list of genes.

**Supplementary Figure 8**

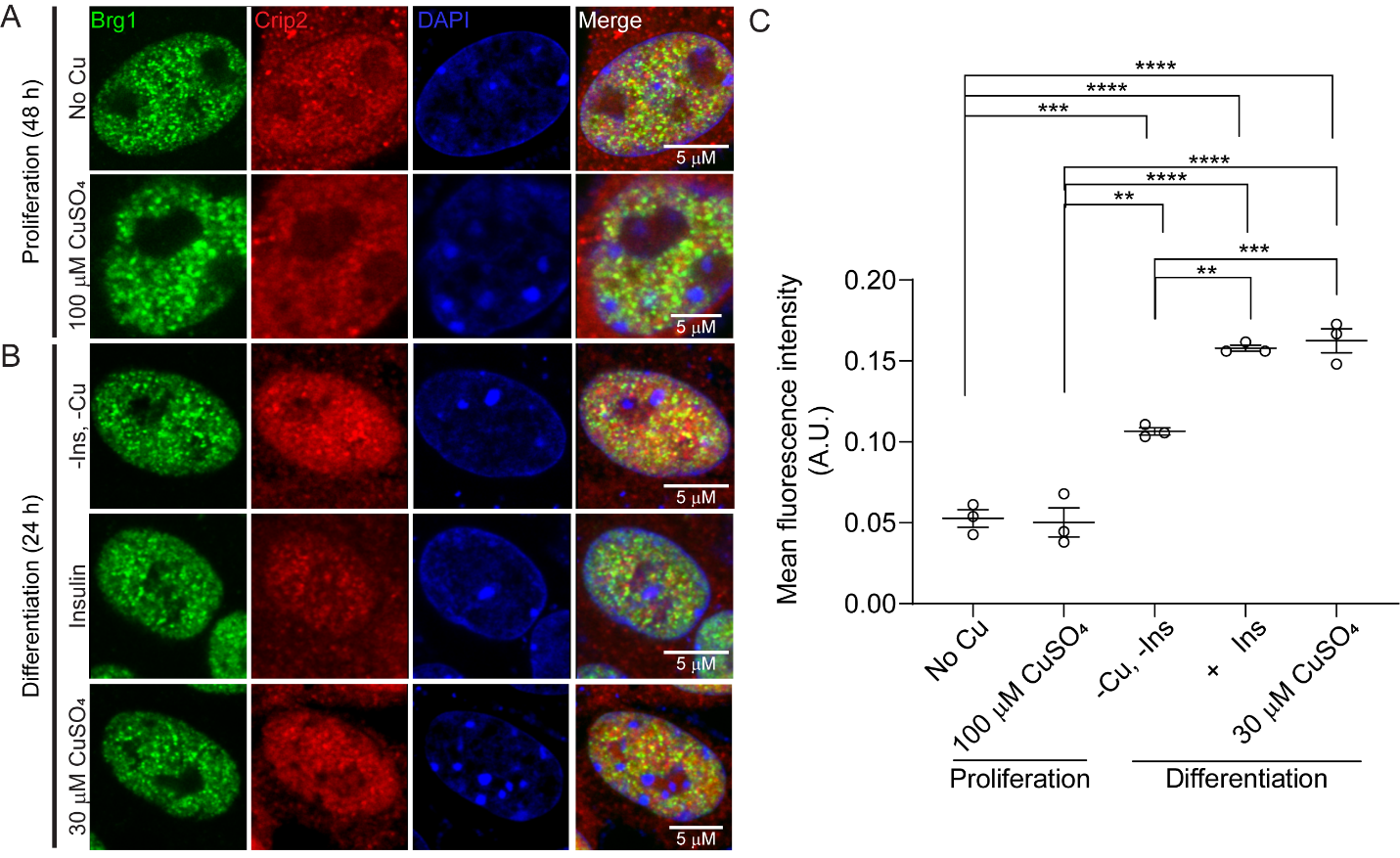

**Supplementary Figure 8. mCrip2 partially co-localizes with Brg1 in primary myoblasts.** Representative confocal microscopy zooming into nuclei stained for the chromatin remodeler enzyme Brg1 (green) and mCrip2 (red) colocalization (merge panel) in proliferating **(A)** and differentiating **(B)** myoblasts supplemented or not with CuSO_4_. The nucleus is stained with DAPI (blue). **C.** Plot represents the quantification of the nuclear colocalization for mCrip2 and Brg1 obtained from at least 20 cells imaged for each biological experiments; N=3. ***P < 0.001; ****P < 0.0001.

**Supplementary Figure 9**

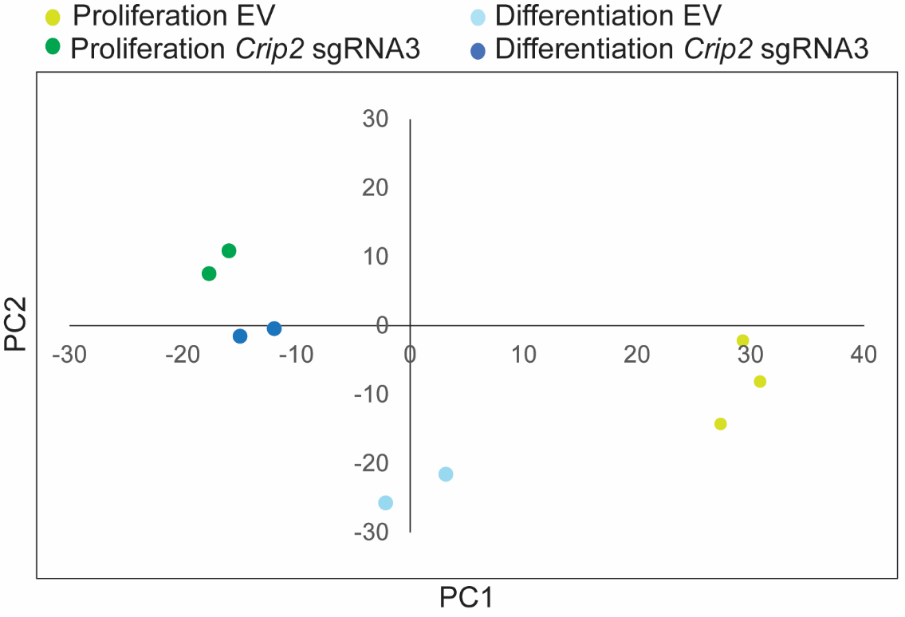

**Supplementary Figure 9. Principal Component Analysis (PCA) plot showing the variance of the control and *Crip2* sgRNA datasets used in the RNAseq analyses.** The PCA plot illustrates the distribution of each RNAseq biological replicate in the reduced-dimensional space defined by the principal components. Control empty vector (EV) proliferating samples are in light green and *Crip2* sgRNA3 samples are in dark green. Control empty vector (EV) differentiating samples are in light blue and *Crip2* sgRNA3 samples are in dark blue.

**Supplementary Figure 10**

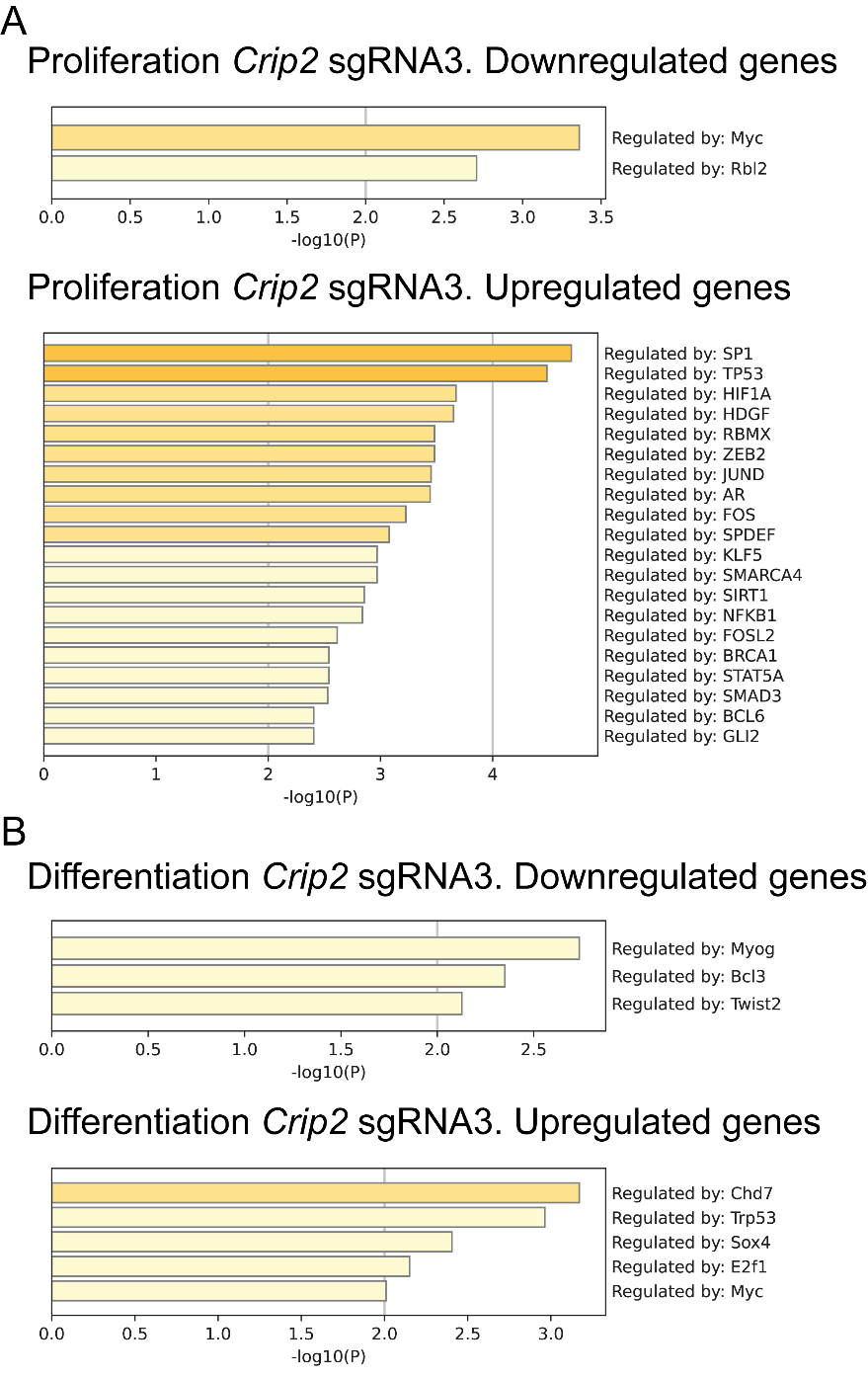

**Supplementary Figure 10.** **TRRUST** **gene set enrichment analysis**. Enrichment plot showing significant enrichment of transcription factors targets among differentially expressed genes in proliferating (**A**) and differentiating (**B**) primary myoblasts depleted of *mCrip2*. The x-axis represents the ranked list of genes based on the significance of differential expression.

**SUPPLEMENTARY FIGURE 11**

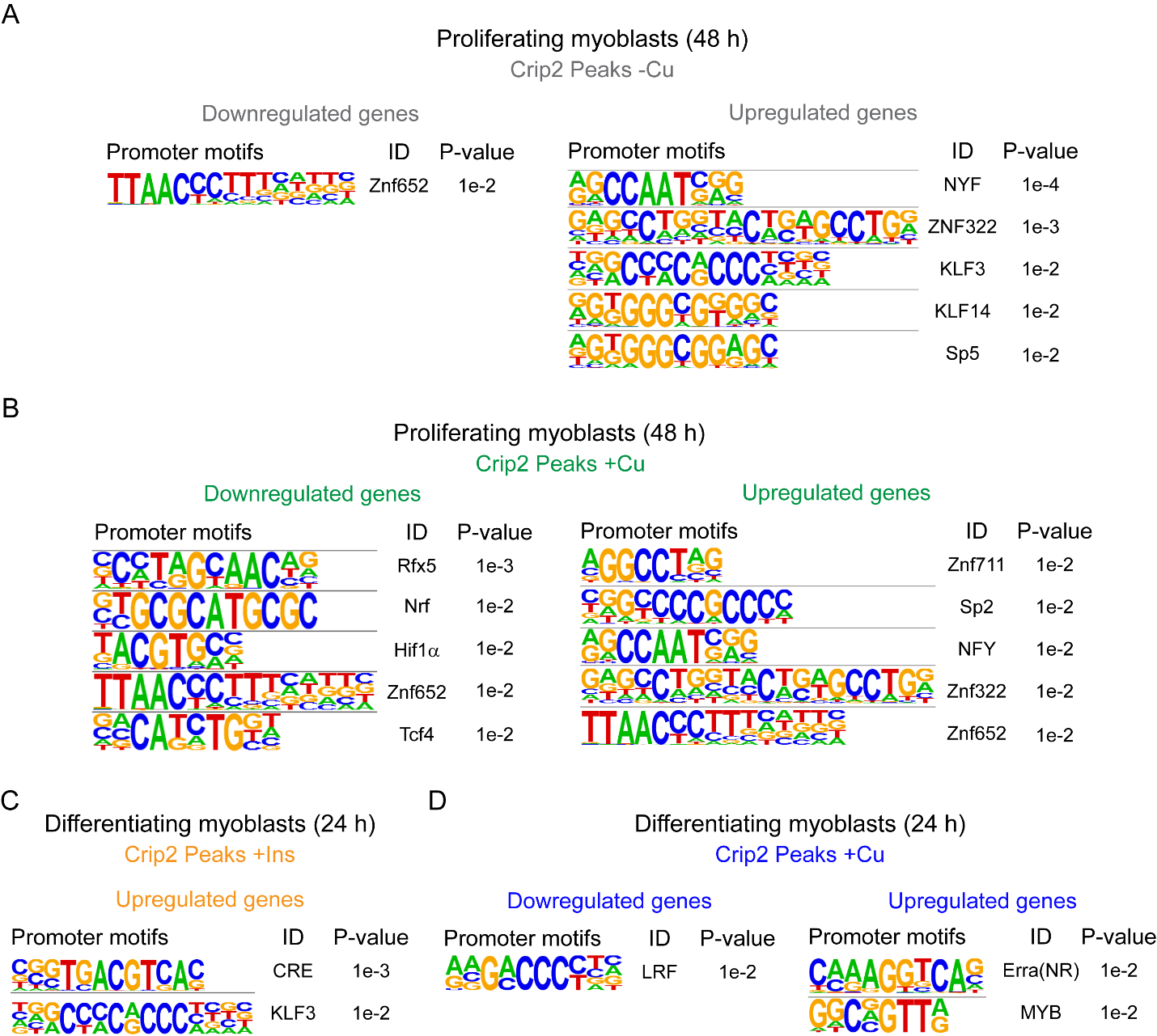

**­­­**

**Supplementary ­­­Figure 11. Main changes of mCrip2 motif-binding dependent on Cu supplementation in proliferating and differentiating primary myoblasts.** Novel consensus DNA-binding motifs identified from peak calling by Sparse Enrichment Analysis (SEACR) for CUT&RUN within mCrip2 peaks in proliferating cells (**A**) and differentiating myoblasts (**B**) supplemented or not with Cu. The top five most significant motifs enriched, including the DNA logo, its corresponding TF, and its P value are shown. See **Supp. table 3** for complete list of peaks. Representative genome browser tracks of CUT&RUN experiments examining mCrip2 binding to the *MyoD1* (**C**), *Mt1* (**D**) and *Mt2* (**E**) promoters in proliferating and differentiating myoblasts (upper panels). *MyoD1, M1 and Mt2* promoters were selected as a representative locus for validation by ChIP-qPCR (lower panels). Plots represent data obtained from 3 independent biological experiments.

**Supplementary Table 2. List of primers used in this study.**

| **Primer Name** | **5’ Sequence** | **3’ sequence** | **Use** |
| --- | --- | --- | --- |
| *Crip2* | GAATTCGCCTCCAAGTGTCCC | CCATGGTTGGGCTGAACTGTGCCTTC | cloning into pR-RIBA and pBABE |
| *Crip2* sgRNA1 | CACCGGGACACTTGGAGGCCATGGT | AAACACCATGGCCTCCAAGTGTCCC | sgRNAs for CRISPR/Cas9 |
| *Crip2* sgRNA2 | CACCGCTTCCTCTCTACAGCTGAGA | AAACTCTCAGCTGTAGAGAGGAAGC | sgRNAs for CRISPR/Cas9 |
| *Crip2* sgRNA3 | CACCGTAGGGCTGGCCATCGTGCTG | AAACCAGCACGATGGCCAGCCCTAC | gRNAs for CRISPR/Cas9 |
| *q-Eef1A* | GGCTTCACTGCTCAGGT  GATTATC | ACACATGGGCTTGCCAGGGAC | Gene  Expression |
| *q-Pax7* | GCAGCTGGAGGAGCTAGAGAAG | GTCTCCTGGCTTGATGGAGTC | Gene  expression |
| *q-Myog* | CAAGTGTGCACATCTGTTCTAGTCTCT | GTATCATCAGCACAGGAGACCTTGGT | Gene  expression |
| *q-Mck* | GCCGGGGATGAGGAGTCCTAC | GCAGTGCGGAGGCAGAGTGTA | Gene  expression |
| *q-Mtf1* | AGATGATATTGTCGTCTGGACTGTG | GAAGCGGAAGTGACGCTAGGGACAG | Gene  expression |
| *q-RpsA* | GGTGGCACCAACCTTGACTTTC | GTCAGCAGGATTCTCGATGGCA | Gene  expression |
| *q-Rps8* | GGAGGCAATAAGAAGTACCGTGC | TTGGTGCGGACAAGCTCGTTGT | Gene  expression |
| *ch-MyoD1* | GCTCAGCAACTATGCTCTACA | CGCCCTCCAAAGCGCACAAAT | ChIP-qPCR |
| *ch-Mt1* | CTCCGCCCGAAAAGTGCGCTC | GAAGCTGGAGCTACGGAGTAA | ChIP-qPCR |
| *ch-Mt2* | TTAGCACACAAGACATGC | TTTCTCCCGAGTCCCTT | ChIP-qPCR |

**Supplementary Table 3.**

**Parameters obtained from Cyclic voltammetry at different scan rates for Au bare and cysteine modified electrodes with hsCRIP2 protein.**

| Protein | E^⸰^  (mV vs Ag/AgCl) | ∆E_p_  (mV) | k_s_  (s^-1^) | α | Γ  (mol^*^cm^-2^) |
| --- | --- | --- | --- | --- | --- |
| ^a^Au-Cys/CRIP2-Cu | 151 | 295 | 1.02 | 0.53 | 1.07x10^-10^ |
| ^a^Au/CRIP2-Cu | n.c | n.c | n.c | n.c | n.c. |
| ^b^Au/Cu-SOD | n.c | n.c | n.c | n.c | n.c. |
| ^b^Au-Cys/Cu-SOD1 | 88 | 213 | 1.05 | 0.54 | 4.28x10^-11^ |

n.c. *not calculated*, due the absence of either the oxidation or reduction signal.

^a^ hsCRIP2 cyclic voltammetry parameters.

^b^ For comparison: Published values obtained for SOD1 by Jimenez-Gonzalez *et al*., 2023
